# USP7 deubiquitinase stabilizes FAN1 to support DNA crosslink repair and suppress CAG repeat expansion

**DOI:** 10.1101/2025.08.08.669300

**Authors:** G Collotta, M Gatti, I Ungureanu, V van Ackeren, E Rannou, F Vivalda, D Gomez Vieito, KM Fishwick, C von Aesch, A Porro, K Ungerleider, A Heidari, R Guérois, RJ Harding, S Bischof, G Balmus, AA Sartori

## Abstract

Human FAN1 is a structure-specific endonuclease critical for the repair of DNA interstrand crosslinks (ICLs) and the excision of extrahelical CAG repeats–whose pathological expansion underlies Huntington’s disease (HD), a progressive and currently incurable neurodegenerative disorder. However, mechanisms of post-translational regulation of FAN1 are still largely unknown. Here, we identify the ubiquitin-specific protease 7 (USP7) as new interactor of FAN1. USP7 stabilizes FAN1 protein levels in a deubiquitination-dependent manner, preventing FAN1 from proteasomal degradation. Consequently, we demonstrate that USP7 depletion leads to reduced chromatin association of FAN1 and increased cellular hypersensitivity following ICL damage. Moreover, we find that loss of USP7 accelerates CAG repeat expansion in an HD cellular model. Collectively, our findings establish USP7 as a critical regulator of FAN1 activity in the maintenance of genome stability, highlighting potential therapeutic opportunities for cancer and HD.

## INTRODUCTION

FAN1 (FANCD2 and FANCI-associated nuclease 1) was originally identified by four independent research groups as a DNA repair enzyme involved in the processing of DNA interstrand crosslinks (ICLs)^1–4^. FAN1 possesses dual catalytic activity, functioning both as a 5′ flap endonuclease and a 5′−3′ exonuclease. This enables FAN1 to unhook ICLs, presumably through a series of precise incisions spaced approximately three nucleotides apart^5–7^. Beyond its role in ICL repair, FAN1 also interacts with mismatch repair (MMR) proteins of the MutL family, including MLH1, PMS2, and MLH3, forming MutLα and MutLγ nuclease complexes^2,3,8^. Notably, MLH1 binding was proposed to restrict FAN1’s nuclease activity to specific DNA substrates, as cells expressing FAN1 mutants defective in MLH1 interaction displayed increased sensitivity to ICL-inducing agents^9,10^. More recently, genome-wide association studies (GWAS) have shown that FAN1 variants are the strongest genetic modifiers in Huntington’s disease (HD), a hereditary and fatal neurodegenerative disorder^11^. Here, FAN1 is thought to compete with the MMR system and suppress somatic expansion of CAG tracts in exon 1 of mutant *Huntingtin* (*mHTT*)^12–16^. Notably, elevated FAN1 expression have been associated with delayed age-of-onset in HD patients, correlating with a slower rate of CAG repeat expansion^17^.

Cellular FAN1 protein levels are relatively low and, typical for enzymes capable of cleaving DNA, its activity expected to be tightly regulated through post-translational modifications, including proteolysis via the ubiquitin-proteasome pathway. The ubiquitination machinery has emerged as a critical player in maintaining genome stability, orchestrating essential DNA damage response (DDR) processes, including multiple DNA repair pathways^18^. Despite this, very little is known about the specific factors that govern FAN1 ubiquitination, aside one report describing APC/C-Cdh1–dependent FAN1 degradation when cells exit mitosis^19^. Protein ubiquitination is a highly dynamic and reversible process, primarily regulated by selective E3 ubiquitin ligases and deubiquitinases (DUBs), which add or remove ubiquitin chains from target proteins, respectively^20^. The human genome encodes around 100 DUBs, categorized into six distinct families. Among these, the ubiquitin-specific protease (USP) family is the largest, with many members being associated with all hallmarks of cancer, most notably genomic instability and mutation^21^.

In this study, using mass spectrometry-based proteomics, we identified the DUB USP7 as a regulator of FAN1. We show that USP7 interacts with FAN1 through its N-terminal TRAF-like and C-terminal ubiquitin-like (UBL) domains. USP7 deubiquitinates FAN1, thereby preventing its proteasomal degradation. Moreover, we demonstrate that USP7 is required for the assembly of FAN1 at damaged chromatin following ICL damage. Loss of USP7 results in elevated apoptotic signalling and hypersensitivity to mitomycin C (MMC). Lastly, we provide evidence that USP7 is necessary for maintaining CAG repeat stability. Collectively, these findings uncover a previously unrecognized mechanism by which USP7 stabilizes FAN1, offering new insights into the post-translational regulation of DNA repair proteins in human cells.

## RESULTS

### Identification of USP7 as an interaction partner of FAN1

To gain insights into the regulation of human FAN1, we interrogated the FAN1 interactome using affinity purification coupled to mass spectrometry (AP-MS). To this end, we established a HEK293 cell line, in which endogenous FAN1 was replaced by an inducible, eGFP-tagged variant, using a stably-integrated doxycycline (Dox)-inducible vector containing two cassettes: one expressing GFP-FAN1 wild-type (wt), and the other an shRNA targeting endogenous *FAN1* mRNA^22,23^. Using this cell line, we performed GFP-Trap co-immunoprecipitation (co-IP) from nuclei-enriched lysates followed by liquid chromatography-tandem mass spectrometry (LC-MS/MS). This analysis recovered known FAN1 interactors, including the MutLα complex (MLH1-PMS2)^8^, the FANCD2/FANCI complex^1–4^, and PCNA^23^ **(Figure 1A and Supplementary Table 1)**. In addition to these established interactors, we also discovered several putative FAN1 regulators, including three USP-type deubiquitinases (USP7, USP9X and USP48), all previously implicated in the DNA damage response^24–27^ **(Figure 1A and Supplementary Table 1)**. USP7 and USP9X have also been detected, but not validated, in an earlier FAN1 AP-MS study^4^.

**Figure 1.**
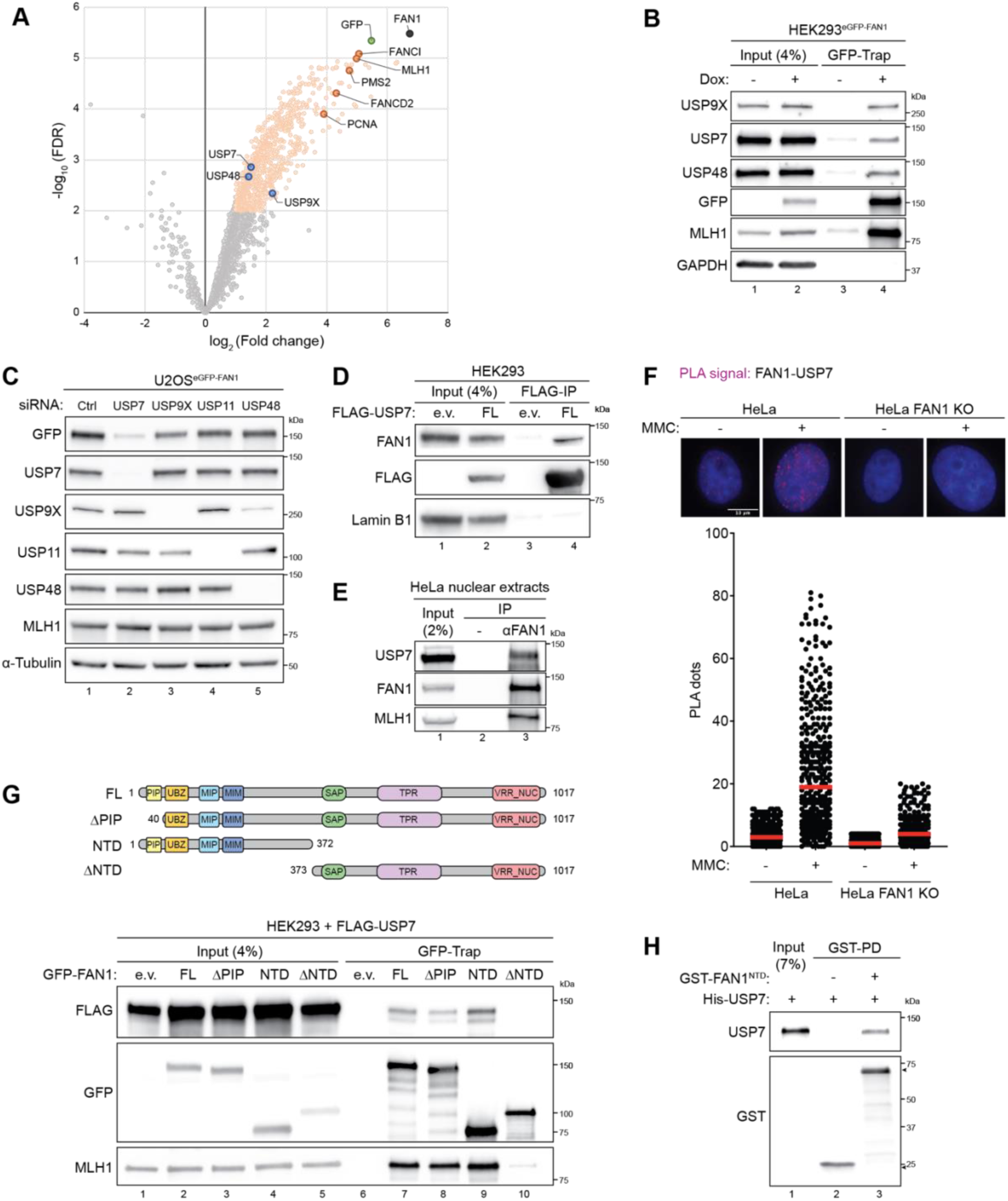
USP7 interacts with the N-terminus of FAN1. (**A**) Volcano plot showing all GFP-FAN1-interacting proteins identified by immunoprecipitation with GFP-Trap beads followed by mass spectrometry in HEK293^eGFP-FAN1^ cells. Significantly enriched proteins (log_2_ (Fold Change) ≥ 1 and –log_10_ (FDR) ≥ 2) are displayed as light orange dots. FAN1, GFP, known FAN1 interactors and DUBs are shown as black, green, dark orange and blue dots respectively. (**B**) HEK293^eGFP-FAN1^ cells were induced with doxycycline (Dox, 100 ng/ml). 48 h later, cells were lysed and whole-cell extracts were subjected to GFP-Trap resin. Inputs and recovered protein complexes were analysed by immunoblotting. (**C**) Doxycycline-inducible U2OS^eGFP-FAN1^ cells were transfected with either non-targeting (Ctrl) or indicated siRNA oligos. 24 h later, GFP-FAN1 expression was induced with Dox (100 ng/ml) for 24 h and whole-cell lysates were analysed by immunoblotting. (**D**) HEK293 cells were transfected with either empty vector (e.v.) or the FLAG-USP7-wt expression constructs. 48 h after transfection, cells were lysed and whole-cell extracts were subjected to IP using anti-FLAG M2 affinity resin. Inputs and recovered protein complexes were analysed by immunoblotting. (**E**) HeLa nuclear extracts were subjected to immunoprecipitation using anti-FAN1 antibody. Inputs and recovered protein complexes were analysed by immunoblotting. (**F**) PLA was used to evaluate FAN1-USP7 association in HeLa parental and FAN1 KO cells, mock-treated or treated with MMC (150 ng/ml) for 24 h. Dot plot shows the number of PLA foci and the median from at least 260 cells for each condition. Representative images are shown. Scale bar: 10 μm. (**G**) Schematic representation of the human FAN1 protein indicating truncation mutants. PIP, proliferating cell nuclear antigen-interacting peptide box; UBZ, ubiquitin-binding zinc-finger domain; MIP, MLH1-interacting protein box; MIM, MLH1-interacting motif; SAP, SAF-A/B, Acinus and PIAS domain; TPR, tetratricopeptide repeat domain; VRR_NUC, virus-type replication-repair nuclease domain; FL, full-length; NTD, N-terminus; ΔNTD, deleted NTD. HEK293 cells were co-transfected with FLAG-USP7 and either e.v. or the indicated GFP-FAN1 expression constructs. 48 h after transfection, cells were lysed and whole-cell extracts were subjected to GFP-Trap resin. Inputs and recovered protein complexes were analysed by immunoblotting. (**H**) Soluble extracts expressing either GST or GST-FAN1-NTD (aa 1-372) were immobilized on Glutathione Sepharose 4B affinity resin and incubated with purified recombinant His-USP7. Inputs and recovered protein complexes were analysed by immunoblotting.

To validate these proteomics data, we performed GFP-Trap pulldowns in HEK293^eGFP-FAN1^ cells and found that endogenous USP7, USP9X, and USP48 co-precipitated with FAN1 **(Figure 1B)**. To assess whether these DUBs affect FAN1 stability, we individually depleted USP7, USP9X and USP48 from U2OS^eGFP-FAN1^ cells. As a control, we included USP11, a DUB not enriched in our AP-MS data set but implicated in DNA repair^28,29^. Remarkably, western blot analysis revealed that silencing of USP7 leads to a significant reduction in GFP-FAN1 levels, while depletion of the other DUBs had no impact on FAN1 stability **(Figure 1C)**. Reciprocal pulldown experiments further revealed that the USP7-FAN1 interaction occurs independently of USP9X **(Figure S1A)**, despite prior reports of USP7-USP9X cooperation in deubiquitinating certain protein substrates^30^.

Given its robust impact on FAN1 protein levels, USP7 was prioritized for further functional investigation. Co-IP of FLAG-USP7 from HEK293 cells confirmed its association with endogenous FAN1 **(Figure 1D)**, and reciprocal co-IP using an anti-FAN1 antibody in nuclear extracts from HeLa cells corroborated this interaction under physiological conditions **(Figure 1E)**. To address whether the USP7-FAN1 interaction is modulated by DNA damage, we performed *in situ* proximity ligation assays (PLA) in HeLa parental and HeLa FAN1 knock-out (KO) cells treated with mitomycin C (MMC). PLA signals significantly increased in numbers upon MMC treatment, suggesting that the FAN1-USP7 association is enhanced following ICL damage **(Figure 1F)**. To map the FAN1 region required for USP7 binding, we performed GFP-Trap pulldowns using FAN1 truncation constructs in HEK293 cells. USP7 associated with the FAN1 N-terminal domain (aa 1-372, NTD), a largely unstructured region of FAN1, while deletion of this region (aa 373-1017, ΔNTD) abolished USP7 binding **(Figure 1G)**. As the FAN1-NTD contains two MLH1-binding motifs^9^, we asked whether USP7-FAN1 interactions depend on MLH1. A synthetic, 60mer FAN1 peptide covering the entire MLH1-binding region (aa 118-177) blocked FAN1-MLH1, but not FAN1-USP7 interaction, indicating that USP7 binds to FAN1 in an MLH1-independent manner **(Figure S1B)**. Consistently, FAN1–USP7 interaction persisted in MLH1-deficient HEK293T cells **(Figure S1C)**. Moreover, treatment with the ubiquitin-activating enzyme (UAE) inhibitor TAK-243^31^, causing depletion of cellular ubiquitin conjugates, did not impact FAN1-USP7 binding, indicating this interaction is largely independent of FAN1 ubiquitination **(Figure S1C)**. Finally, GST-FAN1 pulldown experiments using purified recombinant proteins confirmed a physical interaction between recombinant USP7 and the FAN1-NTD **(Figure 1H)**. Collectively, these findings identify USP7 as a previously uncharacterized FAN1-interacting protein and suggest that USP7 stabilizes FAN1 through direct binding to its N-terminal domain. This interaction is enhanced upon DNA damage and may represent a regulatory axis in the cellular response to genotoxic stress.

### FAN1 associates with both the TRAF-like and UBL domains of USP7

Substrate recognition by USP7, including RNF169, p53 and DNMT1 among many others, is typically mediated through two evolutionarily conserved regions: the N-terminal TRAF-like domain and the tandem UBL1-2 domains at the C-terminus^32–35^. While most substrates preferentially bind one of these domains, a subset—including DNA polymerase iota (Pol ι) and PAF15—has been shown to engage both^36,37^. To assess which USP7 domain interacts with FAN1, we transfected FLAG-tagged full-length (FL) USP7 constructs bearing point mutations in either the TRAF (D164A/W165A) or UBL2 (D762R/D764R) domain^38^ into HEK293^eGFP-FAN1^ or HEK293T cells and performed FLAG-IPs **(Figures 2A and S2A)**. Compared to well-characterized USP7 substrates, FAN1 binding appeared weaker and less sharply defined **(Figures 2A and S2A)**. Nonetheless, the interaction was markedly reduced upon mutation of the TRAF-like domain, indicating that this domain is a principal site of FAN1 recognition, as seen for p53 **(Figures 2A and S2A)**. Structural studies of USP7 binding partners have revealed a conserved P/A/ExxS linear sequence motif as a core recognition element for the TRAF-like domain^33,39,40^. Thus, we conducted *in silico* phylogenetic analysis of mammalian FAN1 homologs and identified four candidate regions matching this motif in the FAN1-NTD **(Figure S2B)**. To functionally assess these, we synthesized biotinylated 12mer FAN1 peptides harbouring individual P/A/ExxS motifs and tested their binding to recombinant USP7 using streptavidin pulldown assays **(Figure 2B)**. Remarkably, only the FAN1 peptide containing the 181-PQSS-184 TRAF-like domain binding motif interacted with USP7 **(Figure 2B)**. Moreover, mutation of the key serine residue 184 to alanine (S184A) abrogated USP7 binding **(Figure 2C)**. Consistently, AlphaFold 3 (AF3)^41^ structural predictions suggest that FAN1-S184 directly contacts the USP7 substrate-binding groove, including residue D164, a critical component of the TRAF-like binding interface **(Figures S2C and S2D)**. However, although FAN1 181-PQSS-184 conforms to a canonical TRAF-like domain binding motif, full-length GFP-FAN1 carrying the S184A mutation retained the ability to co-precipitate with endogenous USP7, suggesting additional or alternative binding interfaces **(Figure 2D)**. Indeed, FLAG–USP7 constructs harbouring the TRAF domain mutation (D164A/W165A) still weakly co-immunoprecipitated FAN1 **(Figure 2A)**, prompting us to explore the role of other USP7 domains in FAN1 binding.

**Figure 2.**
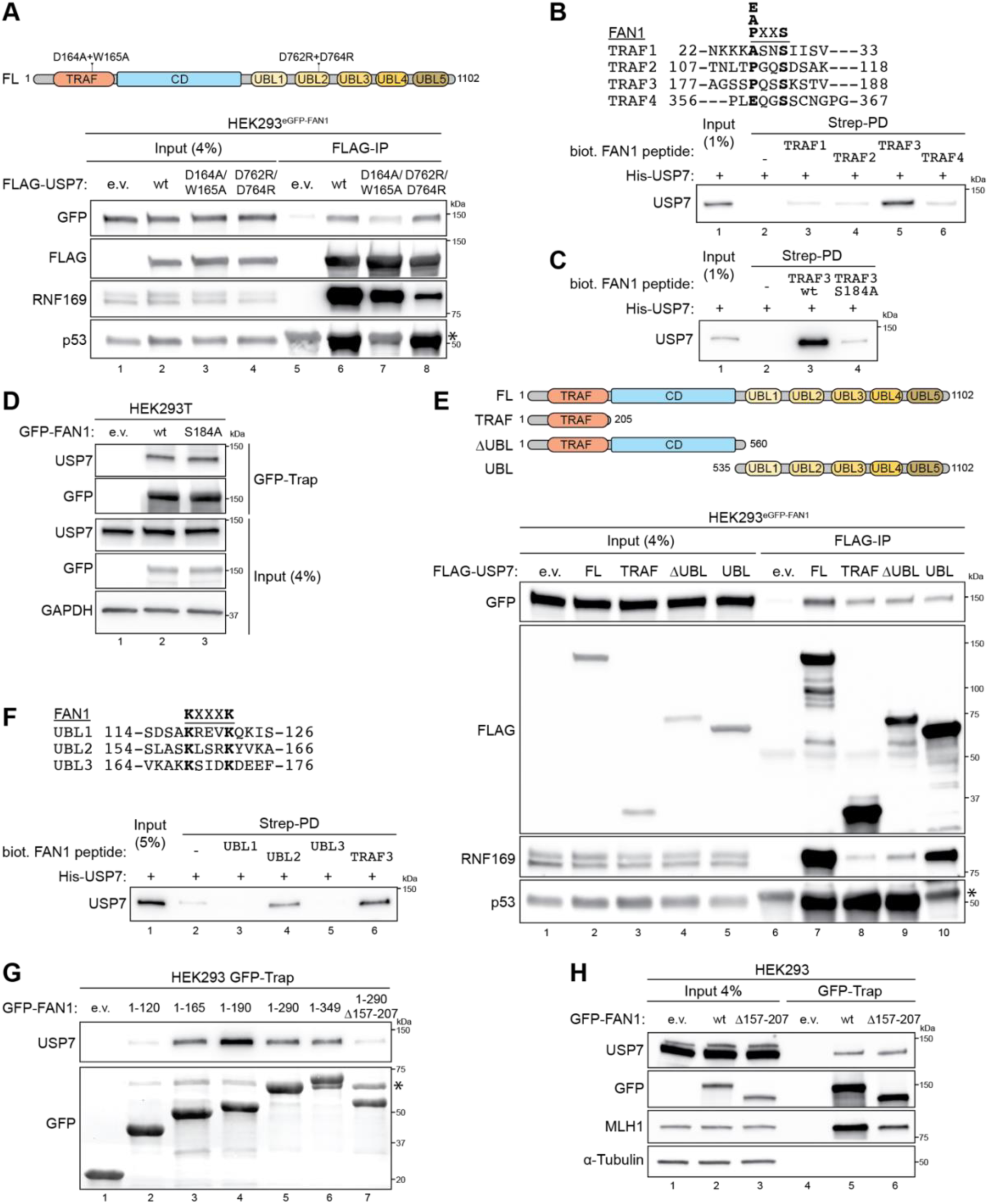
FAN1 associates with the TRAF-like and UBL domains of USP7. (**A**) Top: schematic representation of the human USP7 protein indicating point mutants. Bottom: HEK293^eGFP-FAN1^ cells were transfected with either e.v. or the indicated FLAG-USP7 expression constructs. 24 h later, GFP-FAN1 expression was induced with Dox (100 ng/ml). 48 h post induction, cells were lysed and whole-cell extracts were subjected to IP using anti-FLAG M2 affinity resin. Inputs and recovered protein complexes were analysed by immunoblotting. Asterisk indicates IgG heavy chain from anti-FLAG M2 affinity resin. (**B**) Sequences of the biotinylated 12-mer FAN1-derived TRAF peptides. The corresponding amino acids are indicated. Key residues are highlighted in bold. (**B** and **C**) Biotinylated FAN1 TRAF peptides (10 µg) were immobilized on Strep-Tactin^®^XT-4Flow^®^ resin and incubated with recombinant purified His-USP7. Inputs and recovered His-USP7 were analysed by immunoblotting. (**D**) HEK293T cells were transfected with either e.v. or the indicated GFP-FAN1 expression constructs. 48 h after transfection, cells were lysed and whole-cell extracts were subjected to GFP-Trap resin. Inputs and recovered protein complexes were analysed by immunoblotting. (**E**) Top: schematic representation of the human USP7 protein indicating truncation mutants. TRAF, tumour necrosis factor receptor associated factor; UBL, ubiquitin-like. HEK293^eGFP-FAN1^ cells were transfected with either e.v. or the indicated FLAG-USP7 expression constructs. 24 h later, GFP-FAN1 expression was induced with Dox (100 ng/ml). 48 h post induction, cells were lysed and whole-cell extracts were subjected to IP using anti-FLAG M2 affinity resin. Inputs and recovered protein complexes were analysed by immunoblotting. Asterisk indicates IgG heavy chain from anti-FLAG M2 affinity resin. (**F**) Sequences of the biotinylated 13-mer FAN1-derived UBL peptides. The corresponding amino acids are indicated. Key residues are highlighted in bold. Biotinylated FAN1 UBL peptides (10 µg) were immobilized on Strep-Tactin^®^ XT 4Flow^®^ resin and incubated with recombinant purified His-USP7. Inputs and recovered His-USP7 were analysed by immunoblotting. (**G** and **H**) HEK293 cells were transfected with either e.v. or the indicated GFP-FAN1 expression constructs. 48 h after transfection, cells were lysed and whole-cell extracts were subjected to GFP-Trap resin. Inputs and recovered protein complexes were analysed by immunoblotting. Asterisk indicates unspecific bands.

To that end, we expressed a series of USP7 truncation mutants in HEK293^eGFP-FAN1^ cells and performed FLAG co-IPs. **(Figures 2E)**. In contrast to the domain-specific interactions observed for p53 (TRAF-like) and RNF169 (UBL1-2), GFP-FAN1 co-immunoprecipitated with all USP7 truncation mutants, albeit with reduced efficiency compared to full-length USP7 **(Figure 2E)**. These findings suggest a cooperative contribution of both the TRAF-like and UBL-domains to FAN1 binding. This dual engagement was confirmed using *in vitro* streptavidin pull-down experiments employing recombinant biotin-tagged USP7 TRAF-like or UBL domains, along with MBP-His-tagged FAN1 **(Figure S2E)**. Accordingly, we searched for potential UBL1-2 domain binding sites within the FAN1-NTD that match the consensus KxxxK motif, which has been previously identified in USP7 substrates such as RNF169 and DNMT1^32,42^. The Eukaryotic Linear Motif (ELM) prediction tool revealed five such motifs^43^. Two were excluded due to their location upstream of the FAN1 PIP-box region, which we had previously shown to be dispensable for USP7 interaction **(Figure 1G)**. Streptavidin-pull-down assays with biotinylated FAN1 13mer peptides encompassing the remaining motifs revealed that only the 158-KLSRK-162 sequence bound USP7 **(Figure 2F)**. To validate these findings, we performed GFP-trap pulldown assays using HEK293 cells expressing FAN1 constructs with varying N-terminal lengths. We observed that the FAN1^1–120^ fragment did not interact with USP7, whereas FAN1^1–165^ was the smallest fragment capable of binding to USP7, likely due to the presence of the 158-KLSRK-162 UBL2-binding motif **(Figure 2G)**. USP7 binding was further enhanced in the FAN1^1–190^ fragment, which also includes the 181-PQSS-184 TRAF-like binding motif **(Figure 2G)**. Notably, this enhancement was not observed in longer fragments **(Figure 2G)**. Intriguingly, deletion of a 50-aa region (residues 157-207) from the FAN1^1–290^ fragment, which contains both proposed USP7-binding motifs, substantially disrupted the FAN1-USP7 interaction **(Figure 2G)**. However, targeted deletion of the same 50-aa region from full-length FAN1 did not impair its interaction with USP7 **(Figure 2H)**, suggesting that USP7 may engage alternative proximal TRAF-like or UBL binding motifs on FAN1 in the absence of canonical binding sites, potentially serving as a safeguard to ensure FAN1 regulation. In fact, PAF15 features two P/AxxS and one KxxxK motif, and only a triple mutant completely lost the binding of USP7^37^. Together, these data establish that FAN1 engages both the TRAF-like and UBL1–2 domains of USP7 via distinct linear motifs within its N-terminal region. This bipartite binding mode likely reflects a flexible and robust mechanism of substrate recognition, ensuring dynamic regulation of FAN1 in the context of DNA repair.

### USP7 regulates FAN1 stability through its deubiquitinase activity

Having established USP7 as novel FAN1 interactor, we next sought to dissect its functional impact on FAN1 protein stability. Consistent with our initial observation that USP7 depletion reduced FAN1 levels in U2OS^eGFP-FAN1^ cells **(Figure 1C)**, we investigated this relationship in greater detail using both genetic and pharmacological approaches. We transfected non-transformed, hTERT immortalized retinal pigment epithelial (RPE-1) cells, either parental or p53 KO, with two independent siRNA oligos targeting the coding sequence of *USP7*. In RPE-1 parental p53-proficient cells, USP7 knock-down led to increased p53 protein levels, presumably due to MDM2 degradation^44^, and was accompanied by a pronounced G1 cell-cycle arrest **(Figures 3A and S3A)**. In contrast, p53 KO RPE-1 cells exhibited normal proliferation following USP7 silencing, confirming p53-dependence of the observed cell-cycle effects **(Figures 3A and S3A)**. Notably, FAN1 protein levels were strongly reduced in both cellular backgrounds, paralleling the behaviour of DNMT1, a known USP7 substrate **(Figure 3A)**^34^. To further confirm USP7’s role in FAN1 regulation, we asserted the impact of USP7 inhibition on ectopically expressed FAN1. Knock-down or catalytic inhibition using two non-covalent small molecule inhibitors for USP7^45,46^, led to reduced expression of exogenous FAN1 in Dox-inducible U2OS^eGFP-FAN1^ and HeLa^eGFP-FAN1^ cancer cells **(Figures 3B, 3C and S3B)**. Importantly, GFP-FAN1 levels were restored in USP7-depleted cells upon treatment with the proteasome inhibitor MG132, indicating that USP7 protects FAN1 from proteasome-mediated degradation **(Figure 3D)**. To confirm these results, we examined the FAN1 protein half-life in U2OS^eGFP-FAN1^ cells in the presence of the protein synthesis inhibitor cycloheximide (CHX). FAN1 exhibited a reduced protein half-life upon CHX treatment and this effect was exacerbated by USP7 knock-down **(Figure S3C)**, suggesting that USP7 directly modulates FAN1 protein stability. To determine if USP7’s catalytic activity is required for FAN1 stabilisation, we overexpressed USP7-wt or a catalytic-dead mutant (C223S) in HEK293^eGFP-FAN1^ cells. Only USP7-wt increased GFP-FAN1 levels, while the C223S mutant had no effect **(Figure 3E)**. Consistently, overexpression of USP7-wt significantly reduced FAN1 polyubiquitination in cells co-transfected with HA-tagged ubiquitin, whereas USP7-C223S failed to do so **(Figure 3F)**. To investigate the nature of FAN1 polyubiquitination chains, we employed an HA-ubiquitin mutant retaining only lysine 48 (K48) as acceptor site, with all other lysines mutated to arginine. USP7-wt suppressed K48-linked polyubiquitination of FAN1 whereas the C223S mutant did not, implicating USP7 in the removal of K48-linked chains that signal proteasomal degradation **(Figure 3G)**. Furthermore, when endogenous ubiquitin was the only source for ubiquitination, overexpression of the catalytically-inactive USP7-C223S mutant led to an accumulation of polyubiquitinated FAN1 species, consistent with dominant-negative effect under these conditions **(Figure S3D)**. Finally, we demonstrated that purified recombinant USP7 deubiquitinates immune-purified polyubiquitinated GFP-FAN1 *in vitro* **(Figure 3H)**. Collectively, our results establish USP7 as a bona fide DUB for FAN1. By removing K48-linked polyubiquitination chains, USP7 prevents proteasomal degradation of FAN1 via its enzymatic activity thereby maintaining FAN1 at steady state levels in cells.

**Figure 3.**
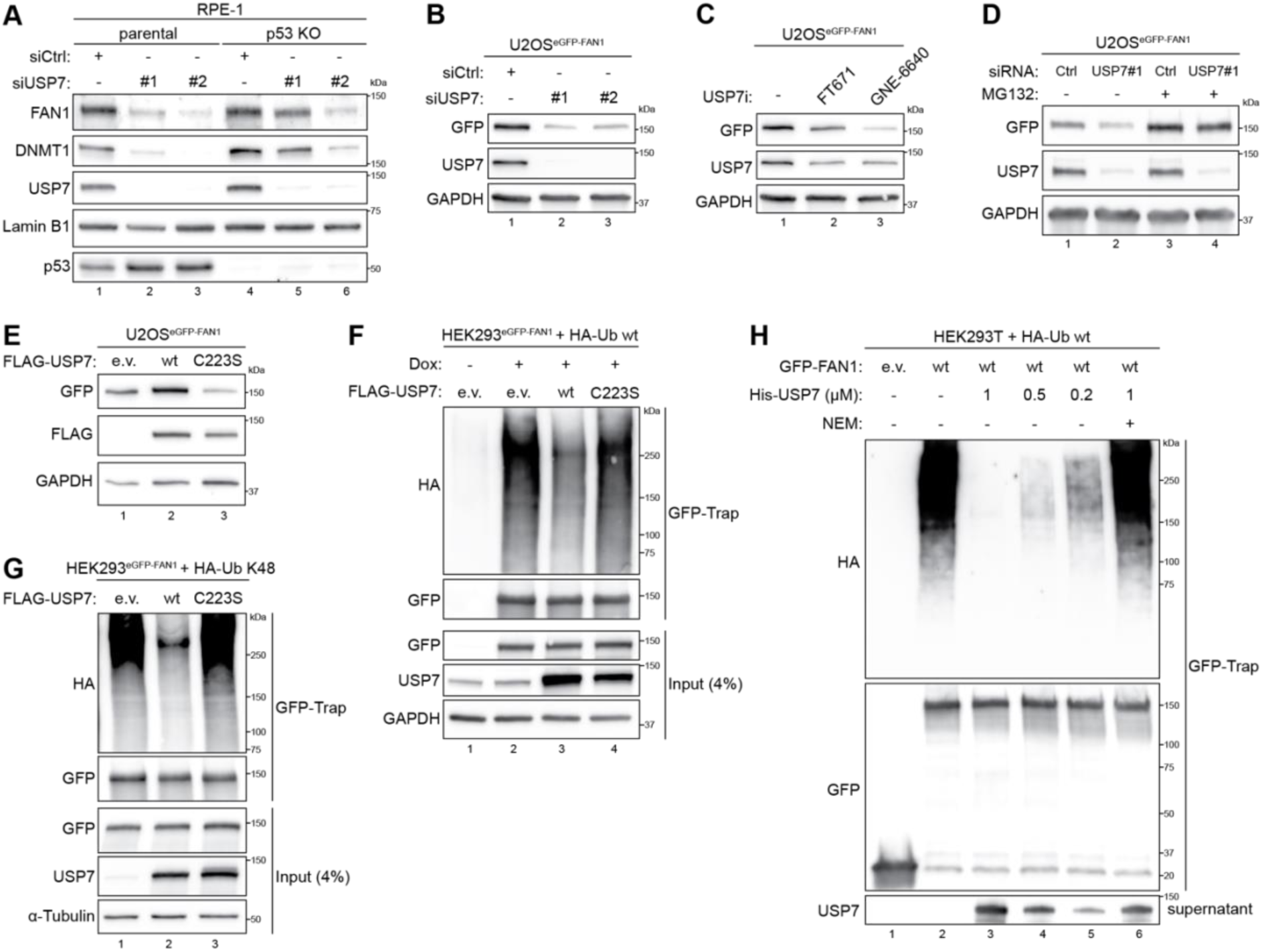
USP7 deubiquitinates FAN1 controlling its protein levels. (**A**) RPE-1 parental and p53 KO cell lines were transfected with either non-targeting (Ctrl) or USP7 siRNA oligos. 48 h later, whole-cell lysates were analysed by immunoblotting. (**B**) Doxycycline-inducible U2OS^eGFP-FAN1^ cells were transfected with either non-targeting (Ctrl) or USP7 siRNA oligos. 24 h later, GFP-FAN1 expression was induced with Dox (100 ng/ml) for 24 h and whole-cell lysates were analysed by immunoblotting. (**C**) Same cells as in **B** were induced with Dox (100 ng/ml). 24 h later, cells were either mock-treated with DMSO (-) or treated (+) with either FT671 or GNE-6640 (10 μM) for 24 h and whole-cell lysates were analysed by immunoblotting. (**D**) Same cells as in **B** were transfected with either non-targeting (Ctrl) or USP7 siRNA oligos. 24 h later, GFP-FAN1 expression was induced with Dox (100 ng/ml). 24 h post induction, cells were either mock-treated (-) or treated (+) with MG-132 (10 μM) for 6 h and whole-cell lysates were analysed by immunoblotting. (**E**) Same cells as in **B** were transfected with either e.v. or indicated FLAG-USP7 expression constructs. 24 h later, GFP-FAN1 expression was induced with Dox (100 ng/ml). 24 h post induction, whole-cell lysates were analysed by immunoblotting. (**F**) Doxycycline-inducible HEK293^eGFP-FAN1^ cells were co-transfected with either e.v. or HA-ubiquitin wt and either e.v. or indicated FLAG-USP7 expression constructs. 24 h later, GFP-FAN1 expression was induced with Dox (100 ng/ml). 24 h post induction, cells were lysed and whole-cell extracts were subjected to GFP-Trap resin. Inputs and recovered protein complexes were analysed by immunoblotting. (**G**) Doxycycline-inducible HEK293^eGFP-FAN1^ cells were co-transfected with either e.v. or HA-ubiquitin K48-only and either e.v. or indicated FLAG-USP7 expression constructs. 24 h later, GFP-FAN1 expression was induced with Dox (100 ng/ml). 24 h post induction, cells were lysed and whole-cell extracts were subjected to GFP-Trap resin. Inputs and recovered protein complexes were analysed by immunoblotting. (**H**) HEK293T cells were co-transfected with HA-ubiquitin wt and either e.v. or GFP-FAN1 expression constructs. 24 h later, cells were lysed and whole-cell extracts were subjected to GFP-Trap resin. Recovered GFP-Trap resin was equilibrated in DUB buffer and then incubated with the indicated concentration of purified recombinant His-USP7 in absence (-) or presence (+) of NEM (20mM).

### USP7 facilitates FAN1 localization and repair activity at ICLs

We earlier showed that MMC-induced DNA damage enhances the interaction between FAN1 and USP7, suggesting that USP7 may primarily function to stabilize or regulate FAN1 activity at sites of ICLs **(Figure 1F)**. To substantiate this possibility using an alternative readout, we scored chromatin-associated FAN1 foci in U2OS^eGFP-FAN1^ cells by using quantitative image-based cytometry (QIBC), a method previously shown to effectively monitor the re-localization of FAN1 in response to genotoxic stress^23^. We found that both siRNA-mediated depletion and catalytic inhibition of USP7 significantly reduced MMC-induced GFP-FAN1 subnuclear foci **(Figures 4A and S4A),** indicating that USP7 is required for efficient retention of FAN1 at ICLs. FAN1 accumulation at sites of ICLs is dependent on FANCD2^47^. However, MMC-induced FANCD2 foci formation remained intact upon USP7 depletion, suggesting that USP7 acts downstream of FANCD2 in modulating FAN1 localization **(Figure 4B)**.

**Figure 4.**
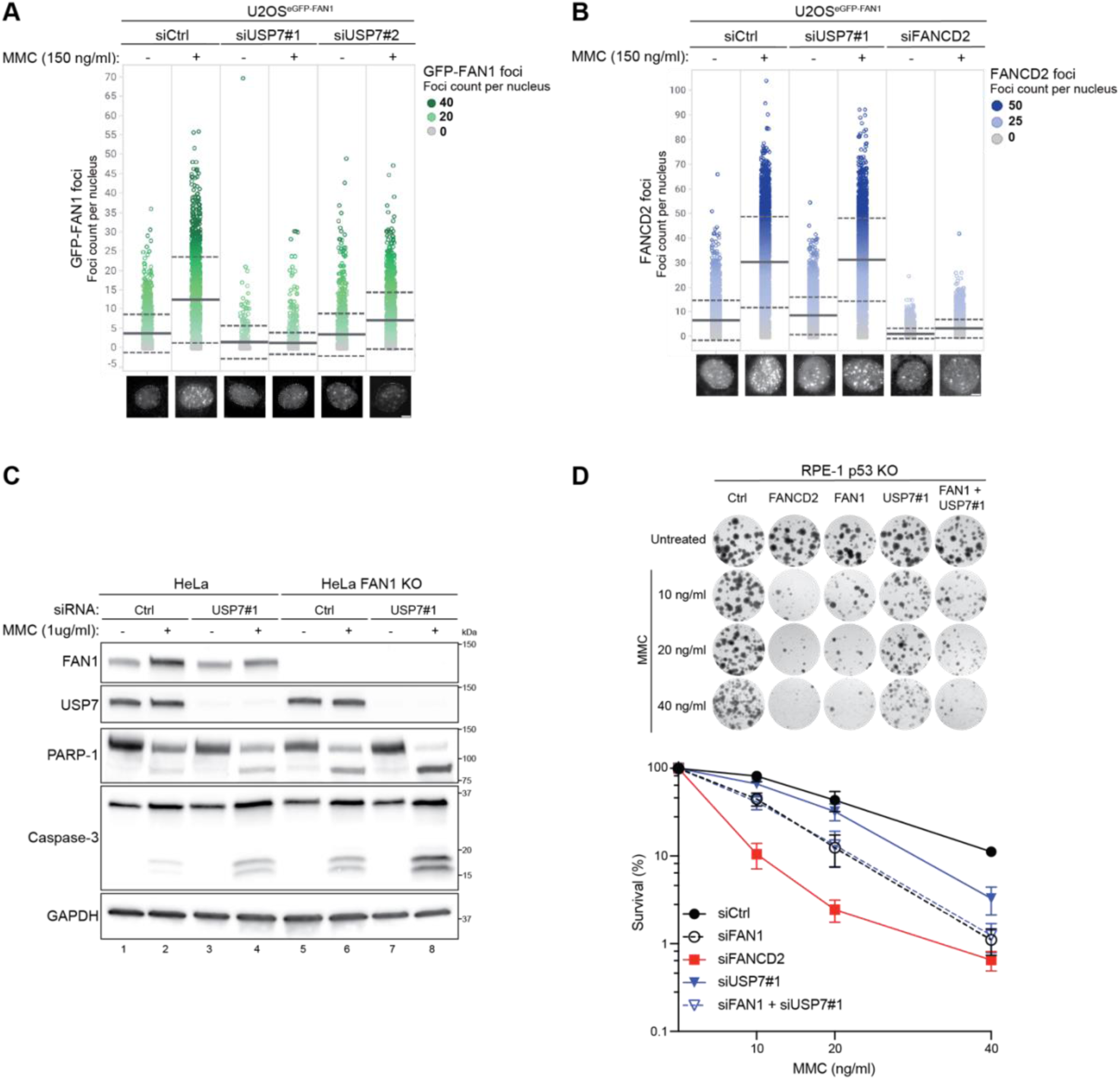
USP7 facilitates FAN1 localization and function during ICL repair. (**A**) QIBC analysis of chromatin-bound GFP-FAN1 foci in U2OS^GFP-FAN1^ cells transfected with either non-targeting (Ctrl) or USP7 siRNA oligos. Cells were either mock-treated (-) or treated (+) with MMC (150 ng/ml) for 24 h. Color-coded scatterplots indicate the number of GFP-FAN1 foci per nucleus. Mean (solid line) and standard deviation (SD) from the mean (dashed lines) are indicated. Representative images are shown below. Scale bar, 10 µm. (**B**) QIBC analysis of endogenous FANCD2 foci in U2OS^GFP-FAN1^ cells transfected with either non-targeting (Ctrl) or the indicated siRNA oligos. Cells were either mock-treated (-) or treated (+) with MMC (150 ng/ml) for 24 h. Color-coded scatterplots indicate the number of FANCD2 foci per nucleus. Mean (solid line) and standard deviation (SD) from the mean (dashed lines) are indicated. Representative images are shown below. Scale bar, 10 µm. (**C**) HeLa parental and FAN1 KO cells were transfected either non-targeting (Ctrl) or USP7 siRNA oligos. 24 h later, cells were either mock-treated (-) or treated (+) with MMC (1 μg/ml) for 48 h and whole-cell lysates were analysed by immunoblotting. (**D**) Clonogenic survival assays of RPE-1 p53 KO cells depleted of the indicated factors and exposed to increasing doses of MMC for 24 h. Viability of untreated cells was defined as 100%. Data are presented as the means ± SD. Representative images are shown (top).

USP7 has been shown to modulate cell survival and apoptosis in response to genotoxic stress, including treatment with DNA crosslinking agents such as cisplatin and MMC^48,49^. We observed that both depletion of USP7 as well as loss of FAN1 in HeLa cells led to increased apoptosis two days after treatment with a high dose of MMC (1 µg/ml), as indicated by elevated proteolytic cleavage of PARP-1 and caspase-3 **(Figure 4C)**. Apoptosis induction was further amplified in HeLa FAN1 KO cells depleted of USP7, suggesting that under these conditions of severe genotoxic stress, USP7 likely regulates multiple factors involved in the repair or signalling of ICL damage^50^ **(Figure 4C)**. These findings were largely recapitulated in MMC-treated RPE-1 p53 KO cells following depletion of FAN1, USP7, or both factors simultaneously, supporting their cooperative role in response to excessive ICL damage **(Figure S4B)**. Notably, FAN1 levels were upregulated in response to MMC, and this induction was abrogated by USP7 depletion, further supporting a role for USP7 in stabilizing FAN1 at ICLs **(Figure 4C)**.

To investigate the functional consequences of USP7 and FAN1 deficiency under sustained genotoxic stress, we conducted clonogenic survival assays following chronic treatment with low doses MMC at 10, 20, and 40 ng/ml. In line with previous reports, depletion of USP7 in HeLa parental cells using two independent siRNA oligos resulted in only a modest increase MMC sensitivity compared to the more pronounced effects observed with RAD18 or FANCD2 depletion^48,49,51^ **(Figures S4C and S4D)**. Notably, USP7 depletion did not exacerbate MMC hypersensitivity in HeLa FAN1 KO cells, in contrast to RAD18 or FANCD2 depletion, which led to an additive increase in sensitivity **(Figures S4E and S4F)**. A similar epistatic relationship was observed in RPE-1 cells: the strongest MMC sensitivity was seen in FANCD2-depleted cells, while FAN1/USP7 co-depletion did not enhance sensitivity beyond individual knock-down of FAN1 **(Figure 4D)**. Collectively, these findings indicate that FAN1 is the primary USP7 substrate involved in the repair of ICLs under moderate DNA damage conditions. However, under high levels of DNA crosslinking stress, the exacerbated apoptotic response in FAN1/USP7 double-depleted cells likely reflects the loss of additional USP7-regulated genome maintenance factors, such as RAD18, RNF168, or SAMHD1^48,49,52^.

### USP7 depletion accelerates CAG repeat expansion

FAN1 is a key modulator of somatic CAG repeat expansion in Huntington’s disease (HD), where its loss leads to increased repeat lengthening—an effect largely driven by the mismatch repair (MMR) pathway, particularly the MutSβ (MSH2–MSH3) complex^53,54^. To establish a robust system to study the somatic expansion of CAG repeats, we generated a human RPE-1 clonal cell model stably expressing *HTT*-exon1 containing 129 CAG repeats (RPE-1^HTT-CAG129^) **(Figures S5A-C)**. In agreement with previous observations in this cellular background^55^, we found that CAG repeat expansion occurred more rapidly in RPE-1^HTT-CAG129^ cells than in U2OS or HD patient-derived induced pluripotent stem cell (iPSC) models **(Figure 5A)**. To validate this model, we performed dual-gRNA CRISPR knock-downs of either FAN1 or MSH3. As expected, FAN1 depletion significantly increased CAG repeat expansion^56,57^, while MSH3 knock-down effectively suppressed repeat instability, consistent with its essential role in driving MMR-dependent expansion^58,59^ **(Figure 5B)**.

**Figure 5.**
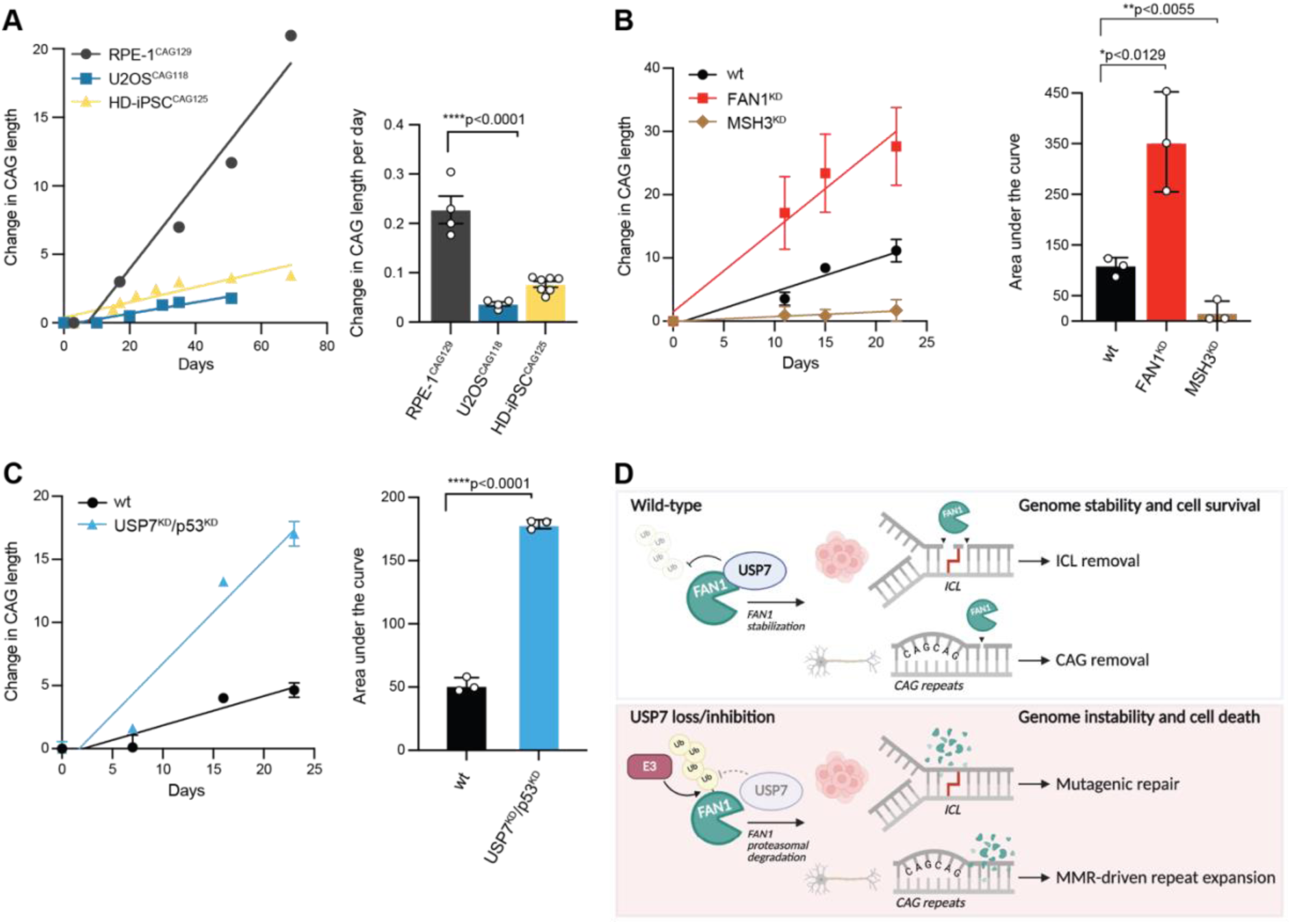
USP7 knock-down accelerates CAG repeat expansion in RPE-1^CAG129^ cells. (**A**) Quantification of CAG repeat expansion across cell lines. CAG lengths were tracked over 80 days in culture. RPE-1^CAG129^ clone C2 exhibited the highest expansion rate compared to HD-iPSC^CAG125^ and U2OS^CAG118^ lines. Bar graph shows daily expansion rates; ****p<0.0001 (one-way ANOVA). (**B**) CAG repeat expansion over time in RPE-1^CAG129^ cells following knock-down of FAN1 (red) or MSH3 (brown) compared to wild-type (wt) controls (black). Left: Line graph showing the mean change in CAG repeat length over 21 days. Right: Quantification of expansion as area under the curve (AUC). Each dot represents a biological replicate (n=3); bars show mean ± SD. Statistical significance was determined by one-way ANOVA with post-hoc multiple comparisons (p = 0.0129 for wt vs. FAN1^KD^; p = 0.0055 for wt vs. MSH3^KD^). This experiment was repeated n=3 independent times. (**C**) Dual knock-down of USP7 and p53 (blue) markedly increases CAG repeat expansion compared to wt cells (black). Left: Time-course analysis reveals a significantly higher expansion rate in double KD cells. Right: AUC quantification confirms the increase (***p < 0.0001, unpaired t-test; n=3). This experiment was repeated n=3 independent times. (**D**) Graphical summary depicting the consequences of USP7 loss, and subsequent FAN1 degradation, for DNA repair. Top: Under physiological conditions FAN1 is subjected to USP7-mediated deubiquitination, preventing FAN1 from being degraded by the proteasome. In human cells, FAN1 is critical for the correct repair of ICLs, where it localizes and unhooks the lesion, rendering the cells resistant to ICL-inducing agents. In neuronal cells, FAN1 is the main nuclease capable of nicking downstream of the CAG repeat, allowing the correct repair of slipped-DNA, therefore delaying HD onset. Bottom: Loss or inhibition of USP7 results in elevated FAN1 polyubiquitination, by a yet unknown E3 ubiquitin ligase, and subsequent proteasomal degradation. Reduction of FAN1 protein levels triggers mutagenic repair of ICLs, which ultimately leads to chromosomal aberrations and cell death. In post-mitotic neurons, lack of FAN1 results in accelerated mismatch repair-dependent CAG repeat expansion, resulting in the accumulation of toxic Huntingtin, driving neuronal loss and disease progression in HD patients.

To determine whether USP7 modulates CAG expansion similarly to FAN1, we used pooled dual-gRNA CRISPR knock-down of USP7 in the RPE-1^HTT-CAG129^ background. To bypass p53-mediated growth suppression following loss of USP7, double knock-down was performed using guides targeting both USP7 as well as p53 (USP7^KD^/p53^KD^). We could confirm effective USP7 depletion **(Figure S5D)**, as well as efficient genome editing **(Figure S5E)**. Moreover, we found that loss of p53 alone (p53^KD^) did not affect CAG expansion relative to the wild-type cells throughout the 35-day analysis **(Figure S5F)**. Strikingly, over a 25-day time course, USP7 knock-down led to significantly accelerated CAG repeat expansion, mirroring the phenotype observed with FAN1 knock-down **(Figures 5B and 5C)**, suggesting that USP7 limits CAG expansion and functions in a pathway akin to FAN1. Together, these data identify USP7 as a novel regulator of CAG repeat stability and demonstrate the strength of the RPE-1^HTT-CAG129^ model for dissecting genetic modifiers of somatic expansion in HD.

## Discussion

FAN1 is a versatile structure-specific DNA endo– and exonuclease involved in several genome stability maintenance pathways, including ICL repair^1,2,4,60^, homologous recombination^1,2^, replication fork integrity^23,61,62^, and, most recently, in the stabilization of expanded trinucleotide repeats^12,13,63^. Despite these diverse roles, very little is known about how the activity of FAN1 is controlled in cells to safeguard genomic integrity. We previously reported that the FAN1-MLH1 interaction is cell-cycle regulated, possibly directing FAN1 nucleolytic activity to specific DNA substrates and preventing otherwise aberrant DNA cleavage^9^. Typical for a DNA repair nuclease, *FAN1* expression levels are low in both cancer and non-cancerous cell lines (<10 RNA transcripts per million, according to v.24 proteinatlas.org)^64^. Therefore, we anticipated that FAN1 protein levels are tightly controlled by the balance of E3 ubiquitin ligases and deubiquitinases. Here, we identify USP7 as a key regulator of FAN1 protein stability. USP7 binds FAN1 via its N-terminal unstructured domain and prevents its proteasomal degradation through deubiquitination. Depletion of USP7 results in MMC hypersensitivity and accelerated rates of CAG repeat expansion, two phenotypes that are associated with FAN1 deficiency.

Mechanistically, we show that USP7 engages FAN1 through a bipartite interaction involving both TRAF-like and UBL1-2 domains. This dual binding mode mirrors previously reported USP7 interactions with substrates such as DNA Polymerase ι, PAF15 and ZMYND8^36,37,65^. Within FAN1, we mapped two putative binding motifs, 181-PQSS-184 and 158-KLSRK-162, that conform to the canonical recognition sequences for USP7. However, mutational analyses suggest the presence of redundant, compensatory or alternative USP7 binding sites in FAN1, consistent with a recent report of USP7 recognizing a non-canonical motif (RxxD) in KRAS^66^. Despite the complexity of this interaction, we found that USP7 robustly removes K48-linked polyubiquitin chains from ubiquitinated FAN1, counteracting FAN1 proteasomal turnover.

Importantly, USP7 was shown to regulate a multitude of cellular processes, including several DNA damage response pathways^50^. Here we could show that USP7 is implicated in ICL repair, at least in part by positively regulating FAN1 activity. Indeed, depletion of USP7 resulted in a reduction in the assembly of FAN1 into subnuclear foci in response to MMC treatment. Moreover, similar to a previous study^67^, we observed that USP7-depleted cells exhibit modest sensitivity to MMC. Importantly, in this context, we found that USP7 and FAN1 function in an epistatic manner, suggesting that FAN1 is a major effector of USP7 in the context of ICL repair.

Genome-wide association studies identified FAN1 as a key genetic modifier of HD onset^12,13^. Subsequently, genetic studies showed that FAN1 deficiency leads to enhanced somatic CAG expansion rates in different cellular and mouse models of HD^68–70^. Currently, there are different models explaining how FAN1 suppresses MMR-driven repeat expansion by competing with the MutSβ complex for the binding and subsequent cleavage of CAG extrusions and/or via its interaction with MLH1^71^. Indeed, using an RPE-1^HTT-CAG129^ – based experimental system, we observed increased CAG repeat instability upon FAN1 inactivation, while repeats are stable over time in MSH3-depleted cells. Remarkably, we found that loss of USP7 phenocopied FAN1 loss, driving CAG expansion at comparable rates. This suggests that USP7 acts as a critical upstream modulator of triplet repeat stability by stabilizing FAN1. Interestingly, USP7 has previously been linked to polyglutamine (polyQ) neurodegenerative diseases, including HD, as it preferentially binds and potentially stabilizes polyQ-expanded mHTT and androgen receptor^72^. Moreover, individuals carrying pathogenic variants in *USP7* are affected by the Hao-Fountain Syndrome (HAFOUS), a neurodevelopmental disorder characterized by global developmental delay, intellectual disability, severe speech delay and behavioural abnormalities^73^. Thus, while our data suggests a potential role of USP7 in preventing somatic repeat expansions in HD brains by stabilizing FAN1 protein levels, we cannot exclude additional functions of USP7 in regulating trinucleotide repeat (TNR) metabolism through additional substrates.

Nevertheless, we believe that our findings could have important therapeutic implications, given that FAN1 protects cells from ICL-induced cell death. In fact, impeding USP7 function could enhance cancer chemosensitivity to DNA crosslinking agents^74^. On the other hand, fostering its activity in stabilizing FAN1 protein levels may have protective effects in a therapeutic setting for carriers of TNR-associated diseases. Indeed, with the same purpose in mind, Cheng and colleagues showed that transfection of codon-optimized *FAN1* mRNA via lipid nanoparticle (LNP) to HD iPSC-derived astrocytes could block CAG repeat expansion. However, delivery of the very same *FAN1* mRNA-LNP to HD mice led to only short duration of increased FAN1 protein expression^75^, highlighting the need for alternative approaches that may include the development of USP7-based deubiquitinase-targeting chimeras (DUBTACs) specific for FAN1^76^.

## ACKNOWLEDGEMENTS

We thank Lorenza Penengo for helpful discussions and providing plasmids for tagged-ubiquitin expression. We thank Sarah Tabrizi and Rob Goold for providing GFP-FAN1 expression constructs. We gratefully acknowledge the Functional Genomics Center Zurich (FGCZ) of University of Zurich and ETH Zurich, for the support on proteomics analyses.

## AUTHOR CONTRIBUTIONS

Conceptualization: G.C., and A.A.S.

Investigation: G.C, M.G., I.U., V.v.A., E.R., F.V., D.G.V., K.M.F., C.v.A, A.P., K.U., and A.H.

Resources: S.B., R.G., R.J.H., and G.B.

Writing – Original Draft: G.C., G.B., and A.A.S.

Writing – Review & Editing: G.C., G.B., and A.A.S.

Supervision, Project Administration & Funding Acquisition: A.P., G.B., and A.A.S.

## MATERIALS AND METHODS

### Cell culture

U2OS, HEK293, HEK293T, HeLa and RPE-1 cells [American Type Culture Collection (ATCC)] were grown in Dulbecco’s modified Eagle’s medium (DMEM) supplemented with 10% fetal calf serum (FCS), penicillin (100 U/mL) and streptomycin (100 μg/mL). U2OS Flp-In T-REx, HEK293 Flp-In T-REx and HeLa Flp-In T-REx (Invitrogen, Life Technologies) cells were maintained in medium supplemented with blasticidin (10 μg/ml) and hygromycin B (250 μg/mL). The Flp-In T-REx system (Thermo Fisher Scientific) was used to generate cell lines stably expressing different short hairpin RNA (shRNA)-resistant forms of eGFP-FAN1 constructs under the control of a doxycycline (Dox)-inducible promoter as previously described^9^. RPE1-Cas9-CAG118 were maintained at 37°C in 5% CO2 and 5% O2. The same cells were cultured in high-glucose DMEM supplemented with 10% fetal bovine serum (FBS), 1% glutamine, 1% non-essential amino acids, penicillin (100 U/mL) and streptomycin (100 μg/mL). HEK293T used for viral production were cultured in high-glucose DMEM, supplemented with 10% fetal bovine serum (FBS), 1% non-essential amino acids, 1% sodium pyruvate and 1% GlutaMAX.

### Virus Production

6-well plates were coated with 0.1% (w/v) gelatine in PBS for 5 minutes. HEK293T cells were seeded at approximately 80% confluency. On day 2, cells were transfected using Lipofectamine™ LTX (Invitrogen) and Opti-MEM™ (Invitrogen). The transfection mixture included the lentiviral transfer vector (targeting USP7, p53, FAN1, or MSH3), packaging plasmid psPAX2, and envelope plasmid pMD2.G, prepared according to the manufacturer’s instructions and incubated for 5 minutes at room temperature after addition of the PLUS reagent. Lipofectamine LTX was then added, and the solution incubated for 30 minutes. Meanwhile, the medium was aspirated from the HEK293T cells, and 5 mL of fresh medium was added. 3 mL of the transfection mixture was added dropwise to each dish. On day 3, the medium was replaced with 8 mL of fresh medium and cells were incubated for an additional 48 hours. On day 6, the virus-containing supernatant was harvested, centrifuged at 1200 rcf for 3 minutes to remove cell debris, and filtered through a 0.45 μm low protein-binding SFCA syringe filter (Nalgene). The virus was aliquoted into 1 mL tubes and stored at –80 °C for up to four months.

### CRISPR-Cas9 gene editing of RPE-1 cells

To generate knock-out cells, RPE1-Cas9-CAG118 cells were seeded at a density of 100,000 cells/mL, and 1:1000 polybrene was added to the mixture. The prepared virus was introduced into the wells at a multiplicity of infection (MOI) of 80% for the following targets: USP7, p53, FAN1 and MSH3. After 3 days, the MOI was assessed using FACS. Cells were selected by treatment with 5 μg/mL puromycin for 3 days. The cells recovered over the course of approximately 7 days. Once the cells reached confluence, samples were collected every 7 days, for a total of 6 time points, with the media being replaced every 4 days.

### Generation of a HeLa FAN1 knock-out cell line

HeLa FAN1 knock-out cells were generated using the CRISPR/Cas9 system. hSpCas9-2A-Puro plasmids (Addgene, nr. 62988) were constructed, each containing the Cas9 protein and a specific guide RNA (gRNA) targeting exon 1 of the *FAN1* gene. The two gRNA target sequences (listed in **Table S2**) were 250 bp apart. Cells were transfected in tandem with both plasmids using Lipofectamine 2000 (Thermo Fisher Scientific) according to the manufacturer’s protocol. Following transfection, cells were selected with puromycin (2 µg/mL) for 48 h to enrich for successfully transfected populations. Individual clones were then isolated and expanded. Clones were screened for CRISPR-mediated genome editing at the target region by sequencing of genomic DNA and knock-out was confirmed by western blot analysis.

### Bacterial strains

Chemo-competent XL1-Blue *E. coli* were used for site-directed mutagenesis and chemo-competent StellarTM cells (TakaraBio) for plasmid cloning using the In-Fusion® technology (TakaraBio). Recombinant GST-FAN1 was expressed in electro-competent BL21-CodonPlus_RIL *E. coli*. Human recombinant full-length USP7 was purchased from R&D Systems (# E-519).

### Bacterial and mammalian expression vectors

All primers used for cloning and site-directed mutagenesis are listed in **Table S3**. Site-directed mutagenesis was carried out by PCR using CloneAmp^TM^ HiFi PCR Premix (TakaraBio) according to the manufacturer’s instructions and subsequent digestion of template DNA with *DpnI* (NEB). pQFlag-USP7 was a gift from Goedele Maertens & Gordon Peters (Addgene # 46751). pcDNA5.1 FRT/TO GFP-FAN1 deletion constructs were kindly shared by Sarah J. Tabrizi (UCL, UK). The PCR product of the USP7 TRAF (residues 2-207) domain was cloned into a pNICBio2 vector containing N-terminal His-tag and C-terminal AviTag through ligation-independent cloning (LIC) using In-Fusion seamless cloning (Takara). The PCR product of the UBL domain of USP7 (residues 520-1067) was cloned into a p28BIOH-LIC vector containing N-terminal AviTag and C-terminal His-tag through LIC using In-Fusion seamless cloning (Takara).

### Purification of recombinant human USP7 fragments

All plasmids were transformed into BL21(DE3)-pBirAcm cells. The cells were grown using a Large-scale EXpression (LEX) bioreactor at 37°C. Once OD600 reached 0.7, the temperature was reduced to 15°C. At OD600 0.8-1, expression of biotinylated proteins was induced by adding 25 mg/L biotin and 0.5 mM isopropyl β-d-1-thiogalactopyranoside (IPTG), and growth continued overnight. Cell cultures were harvested by centrifugation at 7000 rpm, 10 min, 10°C (JLA8.1000). Pellets were flash-frozen in liquid nitrogen and stored at –80°C prior to purification. Cell pellets were thawed, diluted with pre-chilled resuspension buffer (20 mM HEPES pH7.4, 500 mM NaCl, 5% (v/v) glycerol, 1 mM TCEP) and supplemented with 3 μg/ml benzonase, 2 mM MgSO4 and 0.8 mM PMSF/benzamidine-HCl. The mixture was homogenized and lysed by sonication, 60 cycles of 5 s pulse followed by 7.5 s rest. Cell lysates were clarified by centrifugation at 14000 rpm, 1 hr, 10°C (JLA16.250). The supernatant was incubated with TALON resin slurry (2 mL/L pellet) (Takara, #635504) for 30 minutes on an end-to-end shaker at 4°C. Affinity purification was performed using a gravity column. Two wash steps were performed using resuspension buffer, first without and next with 5 mM imidazole. The protein was eluted using 300 mM imidazole. The protein was concentrated using Amicon centrifugal filter units (10 kDa and 50 kDa for TRAF and UBL domains respectively) then loaded onto a HiLoad 16/60 Superdex200 prep grade column (Cytiva) for size-exclusion chromatography on an AKTA pure purification system (Cytiva) using gel filtration buffer (GFB) (20 mM HEPES pH7.4, 300 mM NaCl, 2.5% (v/v) Glycerol, 1 mM TCEP). Peak fractions were concentrated. The protein was flash frozen in liquid nitrogen and stored in –80°C in GFB. Protein purity was analysed by SDS-PAGE and identity was confirmed by mass spectrometry. Biotinylation of proteins were confirmed by streptavidin (Invitrogen, #434301) gel shift assay.

### Purification of recombinant human FAN1

Transformed clones of chemocompetent BL21 bacteria were picked and inoculated in LB medium supplemented with 50 μg/ml kanamycin, subsequently incubated at 37°C overnight under agitation at 250 rpm. The following day, the 5 ml cultures were transferred into a larger Erlenmeyer flask, inoculated again at a 1:50 ratio with LB-medium and shaken at 30°C until the optical density at wavelength of 600 nM (OD600) reached a value between 0.7-0.8. Bacteria were allowed to reach a temperature of 18°C before inducing protein expression by addition of 0.5 mM isopropyl β-D-thiogalactoside (IPTG) (Sigma-Aldrich), with shaking at 18°C and 250 rpm overnight. Bacteria were collected by centrifugation at 4°C and 6000 rpm for 30 min, and the pellet was weighted and resuspended in ice cold lysis buffer (25 mM Tris-HCl pH 8.0, 300 mM NaCl; 1 ml per 4g of pellet). 1 mM PMSF, protease inhibitor cocktail (cOmplete, EDTA-free, Sigma), and 0.1 mg/ml lysozyme were added, and the mixture stirred at 4°C for 30 min. Samples were then sonicated on ice (70% amplitude, 10s on 50s off, 10-minute cycles). 2 μg/ml of Benzonase was added and the samples were incubated rotating at 4°C for 30 min. After a spin-down at 37’000 rpm for 1 hour at 4°C, the insoluble material was removed. Induced proteins were purified from soluble extracts by affinity chromatography using 5 ml HisTrap^TM^ column and proteins eluted using a 20 to 200 mM Imidazole gradient. Elution fractions containing the protein of interest were pooled and purified further by a 5 ml MBPTrap HP column. Elution was performed using a 20 mM maltose elution buffer. Fractions containing the purified protein were additionally run on a HiPrep™ 26/10 Desalting column and purified recombinant full length His6-MBP-FAN1-His6 was stored in 20 mM Tris-HCl pH 8, 150 mM NaCl and 2 mM β-Mercaptoethanol. Purity of the protein was assessed by boiling in SDS sample buffer and separating by SDS-PAGE followed by staining using Coomassie Blue.

### siRNA transfections

Small interfering RNA (siRNA) oligos were purchased from Microsynth and transfected at a final concentration of 40 nM using Lipofectamine RNAiMAX (Thermo Fisher Scientific). A list of all siRNA oligos used throughout this study can be found in **Table S4**.

### Antibodies

A complete list of all primary antibodies used throughout this study can be found in **Table S5**. Secondary horseradish peroxidase (HRP)-conjugated anti-mouse and anti-rabbit antibodies were from GE Healthcare, and the HRP-conjugated anti-goat antibody was purchased from Santa Cruz Biotechnology. Alexa Fluor 488-conjugated secondary antibody was purchased from Thermo Fisher Scientific. The monoclonal antibody against FAN1 was produced by GenScript (NJ, USA) by immunizing mice with recombinant His-tagged FAN1 (aa 871 to 1017) purified from *E. coli*.

### Chemicals and peptides

4′,6-diamidino-2-phenylindole (DAPI), Mitomycin C (MMC), cycloheximide (CHX) and MG132 were purchased from Sigma-Aldrich. FT671 and GNE-6640 were purchased from Aobius. TAK-243 was purchased from MedChemExpress. Doxycycline was purchased from TaKaRa Clontech. A complete list of all peptides used throughout this study can be found in **Table S6**. The synthetic biotinylated FAN1 peptides used in this study were purchased from SynPeptide (Shanghai, China) at 95% purity. Custom-designed FAN1 60-mer peptide (aa 118 to 177) was purchased from GenScript (NJ, USA) and synthesized with a purity >85%.

### Affinity purification coupled to mass spectrometry

HEK293 Flp-In T-REx^eGFP-FAN1^ cells were used to prepare nuclear protein-enriched extracts. Parental HEK293 were used as negative control. After incubation in Swelling Buffer (10 mM Tris-HCl pH 8, 10 mM NaCl) for 10 min at 4°C, cells were collected and centrifuged at 1000 g for 10 min at 4°C. Nuclei were resuspended in IP Buffer (50 mM Tris-HCl pH 7.5, 100 mM NaCl, 1 mM MgCl_2_, 10% Glycerol, 0.1% NP40), supplemented with protease inhibitor cocktail (cOmplete Ultra, EDTA-free, Sigma), treated with Benzonase (100 U) for 15 min at 4°C and disrupted by syringing using a 27G needle. After 21000 g centrifugation for 5 min at 4°C, supernatants containing 1 mg of nuclear enriched proteins were incubated with 25 µL of ChromoTek GFP-Trap® Agarose resin (Proteintech) for 1 hour at 4°C. Beads were washed 4 times in IP buffer, once in IP buffer without NP40 and 4 times in 100 µL of 1X PBS before on-beads tryptic digestion. Mass spectrometry analysis (direct LC/MS-MS) was performed using a high-resolution timsTOF Pro (Bruker) coupled to an Evosep One (Evosep). Samples were separated with the extended Evosep method ‘’15 samples/day’’ keeping the analytical column (PSC-15-100-3-UHPnC, ReproSil C18 3 m 120 Å 15 cm ID 100 µm, PepSep) at 50 °C. For the dual timsTOF, MS spectra were scanned from m/z 100 to m/z 1700 in ddaPASEF mode (data dependent acquisition Parallel Accumulation Serial Fragmentation). For the ion mobility settings, the inversed mobilities from 1/K0 0.60 Vs/cm2 to 1.60 Vs/cm2 were analysed with ion accumulation and ramp time of 166 ms, respectively. 1 survey TIMS-MS scan is followed by 10 PASEF ramps for MS/MS acquisition, resulting in a 1.9 s cycle time. Singly charged ions are excluded using the polygon filter mask and isolation windows for MS/MS were set to m/z 2.0 for precursor ions below m/z 700, and m/z 3.0 for ions above. Then, the acquired MS data were processed using the Fragpipe v17 (MSFragger search engine). The spectra were searched using the Homo sapiens (UP000005640) database with the variable modifications Acetyl (Protein N-term) and Oxidation (M), and the fixed modification Carbamidomethyl (C). Proteomics differential expression analysis was performed with the R-package prolfqua using default settings^77^. Raw data were deposited in the PRIDE database, with the dataset identifier PXD065101.

### Pull-down assays

For GST pull-down assays, GST fusion plasmids were transformed in BL21 RIL (CodonPlus) *E. coli* (Stratagene) and recombinant proteins were expressed by incubating the bacteria for 24 h at 16 °C after the addition of 100 μM IPTG. After centrifugation, the bacterial pellet was resuspended in cold PBS, supplemented with 1% Triton X-100 and protease inhibitors (1 mM PMSF, 1 mM benzamidine, and cOmplete™ Protease Inhibitor Cocktail). After sonication and centrifugation, GST-tagged FAN1 was purified from soluble extracts using Glutathione Sepharose 4 Fast Flow beads (GE Healthcare). GST fusion proteins bound to glutathione beads were mixed with 0.5 µg of human recombinant USP7 (R&D Systems) and incubated for 1 h at 4 °C in 1 ml of PBS-1% Triton. Beads were then washed three times with NTEN300 buffer (0.5% NP-40, 0.1 mM EDTA, 20 mM Tris-HCl pH 7.4 and 300 mM NaCl) and once with TEN100 (20 mM Tris-HCl pH 7.4, 0.1 mM EDTA and 100 mM NaCl) buffer. Recovered complexes were boiled in SDS sample buffer and analysed by SDS–PAGE followed by immunoblotting. For peptide pull-down assays, biotinylated FAN1 peptides were incubated with Strep-Tactin^®^ XT 4Flow^®^ resin (IBA LifeScience) at room temperature for 30 minutes and washed four times with 0.1% Triton-PBS and two times with HNE buffer (20 mM HEPES pH 7.5, 0.2 mM EDTA, 0.5% NP-40, 150 mM NaCl, 0.5 mM DTT, 0.1% BSA, 0.5 mM PMSF). Beads were then mixed with 0.5 µg of human recombinant USP7 (R&D Systems) and incubated rotating for 2 h at 4 °C. Beads were then washed three times with HNE buffer, once with TEN300 (20 mM Tris-HCl pH 7.4, 0.1 mM EDTA and 300 mM NaCl) and once with TEN100 buffer. Complexes were boiled in SDS sample buffer and analysed by SDS–PAGE followed by immunoblotting.

### Co-immunoprecipitation

Cells were lysed in NP-40 buffer [50 mM tris-HCl (pH 7.5), 120 mM NaCl, 1 mM EDTA, 6 mM EGTA, 15 mM sodium pyrophosphate, and 1% NP-40, supplemented with phosphatase inhibitors (20 mM NaF and 1 mM Na_3_VO_4_) and protease inhibitors (1 mM benzamidine, 0.1 mM PMSF and cOmplete™ Protease Inhibitor Cocktail)], incubated with Benzonase (Merck) for at least 30 min at 4°C, and clarified by centrifugation. Between 1 and 3 mg of lysates were incubated with anti-FLAG M2 (Sigma-Aldrich) or ChromoTek GFP-Trap® Agarose resin (Proteintech) for 2 h or overnight at 4°C. The beads were then washed three times with NP-40 buffer and once with TEN100 buffer. Retrieved protein complexes were boiled in SDS sample buffer and analysed by SDS-PAGE followed by immunoblotting.

### FAN1 immunoprecipitation

5 mg of HeLa nuclear extract were pre-cleared for 2 hours at 4°C with rotation. Two micrograms of anti-FAN1 rabbit polyclonal antibody were added to the sample and incubated overnight at 4°C with rotation. Protein A Sepharose beads (CL4B, Sigma-Aldrich) were equilibrated in HeLa nuclear extracts buffer [20 mM HEPES (pH 7.9), 100 mM KCl, 0.2 mM EDTA, 20% Glycerol, 0.5 mM PMSF, 0.5 mM DTT], and 30 μl of bead slurry was added to each sample and incubated for 2 hours at 4°C with rotation. The beads were washed three times with HeLa nuclear extracts buffer and once with TEN100 buffer. Retrieved protein complexes were boiled in SDS sample buffer and analysed by SDS-PAGE followed by immunoblotting.

### *In vivo* ubiquitination assay

HEK293 Flp-In T-REx^eGFP-FAN1^ cells were co-transfected with HA-ubiquitin and FLAG-USP7 expression constructs, respectively. 48 h after transfection, cells were scraped in 500 μl of RIPA buffer (50 mM Tris-HCl, pH 7.5, 150 mM NaCl, 1% IGEPAL, 0.5% sodium deoxycholate, 0.1% SDS) supplemented with phosphatase inhibitors (20 mM NaF and 1 mM Na_3_VO_4_), protease inhibitors (1 mM benzamidine and 0.1 mM PMSF) and the deubiquitinases inhibitor N-ethylmaleimide (NEM, 20 mM). Between 1 and 3 mg of lysates were incubated with ChromoTek GFP-Trap® Agarose resin (Proteintech) for 2 h or overnight at 4°C. The beads were then washed once with RIPA buffer, two times with NTEN 500 (0.5% NP-40, 0.1 mM EDTA, 20 mM Tris-HCl pH 7.4 and 500 mM NaCl), once with TEN300 and once with TEN100 buffer. Retrieved protein complexes were boiled in SDS sample buffer and analysed by SDS-PAGE followed by immunoblotting.

### *In vitro* deubiquitination assay

HEK293T cells were co-transfected with plasmids encoding HA-ubiquitin wt and GFP-FAN1 wt, respectively. 24 h after transfection, cells were washed twice with cold PBS and directly lysed in 1 ml of RIPA buffer (50 mM Tris-HCl pH 7.5, 150 mM NaCl, 1% IGEPAL, 0.5% sodium deoxycholate, 0.1% SDS) supplemented with 2mM MgCl_2_, cOmplete inhibitor cocktail (Roche), phosphoSTOP (Roche), 25U/ml benzonase, 0.1 mM PMSF and 20mM N-ethylmaleimide (NEM). Lysates were incubated for 5 min at room temperature and then centrifuged at 15’000 rcf for 15 min. Between 1 and 3 mg of lysates were incubated with ChromoTek GFP-Trap® Agarose resin (Proteintech) for 2 h or overnight at 4°C. Beads were then collected by centrifugation, washed once with RIPA buffer, two times with NTEN 500 (0.5% NP-40, 0.1 mM EDTA, 20 mM Tris-HCl pH 7.4 and 500 mM NaCl), once with TEN300 and four times with TEN100 buffer. Beads were then equilibrated in DUB buffer (50 mM Tris-HCl, pH 7.5, 50 mM NaCl, 1 mM MgCl_2_, 1mM DTT), equally distributed in reaction tubes and incubated with purified recombinant His-USP7 for 2 h at 37°C at low rpm. The unbound fraction (supernatant) was separated from the affinity resin following centrifugation, and the reaction was blocked by the addition of 5x SDS sample buffer. Samples were boiled and analysed by SDS-PAGE followed by immunoblotting.

### Immunoblotting

If not stated otherwise, cells were lysed in RIPA buffer (50 mM Tris-HCl, pH 7.5, 150 mM NaCl, 1% IGEPAL, 0.5% sodium deoxycholate, 0.1% SDS) supplemented with phosphatase inhibitors (20 mM NaF and 1 mM Na_3_VO_4_) and protease inhibitors (1 mM benzamidine and 0.1 mM PMSF), incubated with Benzonase (Merck) for at least 30 min at 4°C, and clarified by centrifugation. Proteins were resolved by SDS-PAGE and transferred to nitrocellulose membranes. Immunoblots were performed with the indicated antibodies, and proteins were visualized using the Advansta WesternBright ECL reagent and the Fusion Solo S imaging system.

### Colony formation assays

HeLa and RPE-1 p53 KO cells were transfected with the indicated siRNA. The next day, the cells were seeded in six-well plates. The next day, the cells were treated with MMC for 24 h. Plates were then washed twice with 2 ml of PBS before adding fresh growth medium to the cells, that were then cultured for 13 days at 37°C before fixation with crystal violet solution [0.5% crystal violet and 20% ethanol (w/v)]. For analysis, plates were scanned and analysed with the ImageJ Plugin ColonyArea using the parameter “intensity percent”.

### Flow cytometry analysis

RPE-1 parental and p53 KO cells were transfected with the indicated siRNA. 48h later, cells were incubated with 10 µM 5-Ethynyl-2′-deoxyuridine (EdU) for 30 min at 37°C. Cells were then harvested by trypsinization, washed, and fixed in 4% Formaldehyde in PBS. EdU labelling was carried out using the Click-iT EdU technology (Thermo Fisher Scientific) as described in the manufacturer’s instructions. DNA was stained by incubating the cells in 1 % BSA/PBS containing 0.1 mg/ml RNase and 1 μg/ml DAPI. A minimum of 20,000 events were recorded with an Attune NxT flow cytometer.

### Immunofluorescence microscopy

For high-content microscopy and QIBC analyses, cells were seeded on sterile 12 mm glass coverslips and allowed to proliferate until they reached a cell density of 70 – 90%. Cells were then washed once with PBS before fixation in 3% formaldehyde in PBS for 15 min at room temperature, washed once in PBS, permeabilized for 5 min at room temperature in 0.2% Triton X-100 in PBS, washed twice in PBS, and incubated in blocking solution (filtered DMEM containing 10% FBS and 0.02% Sodium Azide) for 15 min at room temperature. When the staining was combined with an EdU Click-iT reaction, this reaction was performed prior the incubation with primary antibody according to the manufacturer’s recommendations (Thermo Fisher Scientific). Where indicated, cells were pre-extracted in 0.2% Triton X-100 in PBS for two minutes on ice prior fixation. All primary antibodies were diluted in blocking solution and incubated for 2 h at room temperature. Secondary antibodies (Alexa Fluor 488, 568, 647 anti-mouse and anti-rabbit IgG from Thermo Fisher Scientific) were diluted 1:500 in blocking solution and incubated at room temperature for 1 h. Cells were washed once with PBS and incubated for 10 min with 4’,6-Diamidino-2-Phenylindole Dihydrochloride (DAPI, 0.5 mg/ml) in PBS at room temperature. Following three washing steps in PBS, coverslips were briefly washed with distilled water and mounted on 5 ml Mowiol-based mounting media (Mowiol 4.88 (Calbiochem) in Glycerol/TRIS).

### Quantitative image-based cytometry (QIBC)

Automated multichannel wide-field microscopy for QIBC was performed using the Olympus ScanR System as described previously^78^. Images of cell populations were acquired under non-saturating conditions with Olympus ScanR Image Acquisition software 3.2 and 3.3.0, typically 16 (4×4) images per coverslip, and identical settings were applied to all samples of the same experiment. Images were analysed with the inbuilt Olympus ScanR Image Analysis Software Version 3.2 and 3.3.0, a dynamic background correction was applied, and nuclei segmentation was performed using an integrated intensity-based object detection module based on the DAPI signal. All downstream analyses were focused on properly detected nuclei containing a 2C-4C DNA content as measured by total and mean DAPI intensities. Fluorescence intensities were quantified and are depicted as arbitrary units. Color-coded scatterplots of asynchronous cell populations were generated with Spotfire data visualization software (TIBCO Spotfire 10.10.1.7). Within one experiment, similar cell numbers were compared for the different conditions. For visualizing discrete data in scatterplots, mild jittering (random displacement of data points along discrete data axes) was applied to demerge overlapping data points. Representative scatterplots and quantifications of independent experiments, typically containing several thousand cells each, are shown.

### *In Situ* Proximity Ligation Assay

HeLa parental and FAN1 KO cells were grown in the absence or presence of MMC (150 ng/ml) for 24 hours. Cells were pre-extracted with CSK buffer containing 0.5% of Triton™ X-100 (Sigma–Aldrich) for 5 min on ice and fixed in 4% formaldehyde in PBS (w/v) for 20 min at room temperature (RT). Coverslips were then washed with PBS and stored overnight at 4°C. In situ PLA was performed using Duolink PLA technology (Sigma-Aldrich) according to the manufacturer’s instructions. In brief, coverslips were blocked for 30 min at 37°C with blocking solution and then incubated with the respective primary antibodies for 2 h at 37°C. Coverslips were washed three times for 5 min in Wash Buffer A (0.01 M Tris, 0.15 M NaCl, and 0.05% Tween 20). Then, Duolink anti-Mouse PLUS and anti-Rabbit MINUS PLA probes were coupled to the primary antibodies for 1 h at 37°C. After three wash steps in Wash buffer A for 5 min, the PLA probes were ligated for 30 min at 37°C. Coverslips were then washed three times for 5 min in Wash buffer A. Amplification using the “Duolink In Situ Detection Reagents FarRed” (Sigma-Aldrich) was performed at 37°C for 100 min. After amplification, coverslips were washed twice in Wash Buffer B (0.2 M Tris and 0.1 M NaCl) for 10 min and incubated for 30 min at 37°C with the appropriate secondary antibodies. Coverslips were then washed twice with Wash Buffer B and once in 0.01× Wash Buffer B for 1 min. Lastly, the coverslips were mounted using Vectashield Mounting Media (Vector Laboratories) containing DAPI, sealed and imaged on a Leica DMI 6000 fluorescence microscope at x63 magnification. Analysis of PLA signals was performed using CellProfiler™.

### DNA Extraction

Genomic DNA was isolated from each sample using the DNeasy Blood & Tissue Kit (QIAGEN). The expanded CAG tract was then amplified by PCR, using 150 ng of template DNA and the following primers: 6-FAM–labelled forward primer HD3F and reverse primer HD5. Thermocycling conditions were 95 °C for 10 min; 30 cycles of 95 °C for 30 s, 58 °C for 30 s and 72 °C for 90 s; followed by a final extension at 72 °C for 7 min. PCR products were then run on an ABI 3730xl Genetic Analyzer (Thermo Fisher Scientific) with the MapMarker 1000 ROX size standard (Tebubio).

GeneMapper (Thermo Fisher Scientific) was then used for the alignment of the size standard for all samples. The median, standard deviation and the instability index were calculated using the, a custom program (Romeo package) available at https://michaelflower.org.

