## Supplementary Information for "USP7 deubiquitinase stabilizes FAN1 to support DNA crosslink repair and suppress CAG repeat expansion"

**Supplementary Figures S1 to S5.**

**Supplementary Tables 2 to 6.**

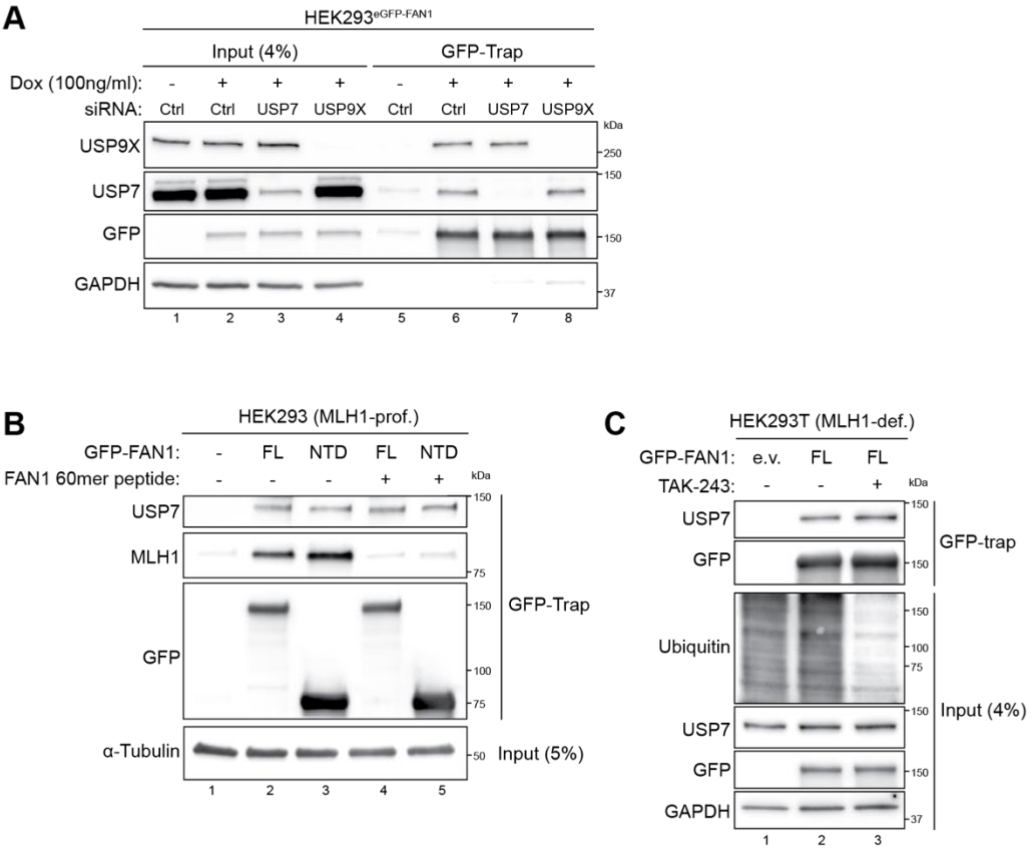


**Figure S1 (related to Figure 1). USP7 associate with FAN1 independently of USP9X, MLH1 or ubiquitin.**

(**A**) HEK293$^{eGFP-FAN1}$ cells were transfected either with non-targeting (Ctrl) or the indicated siRNA oligos. 24 h later, GFP-FAN1 expression was induced with Dox (100 ng/ml). 24 h post induction, cells were lysed and whole-cell extracts were subjected to GFP-Trap resin. Inputs and recovered protein complexes were analysed by immunoblotting. (**B**) HEK293 cells were transfected with either e.v. or GFP-FAN1-wt expressing constructs. 48 h later, cells were lysed and whole-cell extracts were incubated with a FAN1-derived 60-mer peptide (aa 118 to 177) for 2 h and subjected to GFP-Trap resin. Inputs and recovered protein complexes were analysed by immunoblotting. (**C**) HEK293T cells were transfected with either e.v. or GFP-FAN1-wt expressing constructs. 48 h later, cells were either mock-treated (-) or treated (+) with TAK-243 (10 µM) for 2 h, lysed and whole-cell extracts were subjected to GFP-Trap resin. Inputs and recovered protein complexes were analysed by immunoblotting. Molecular weight is indicated on the right.

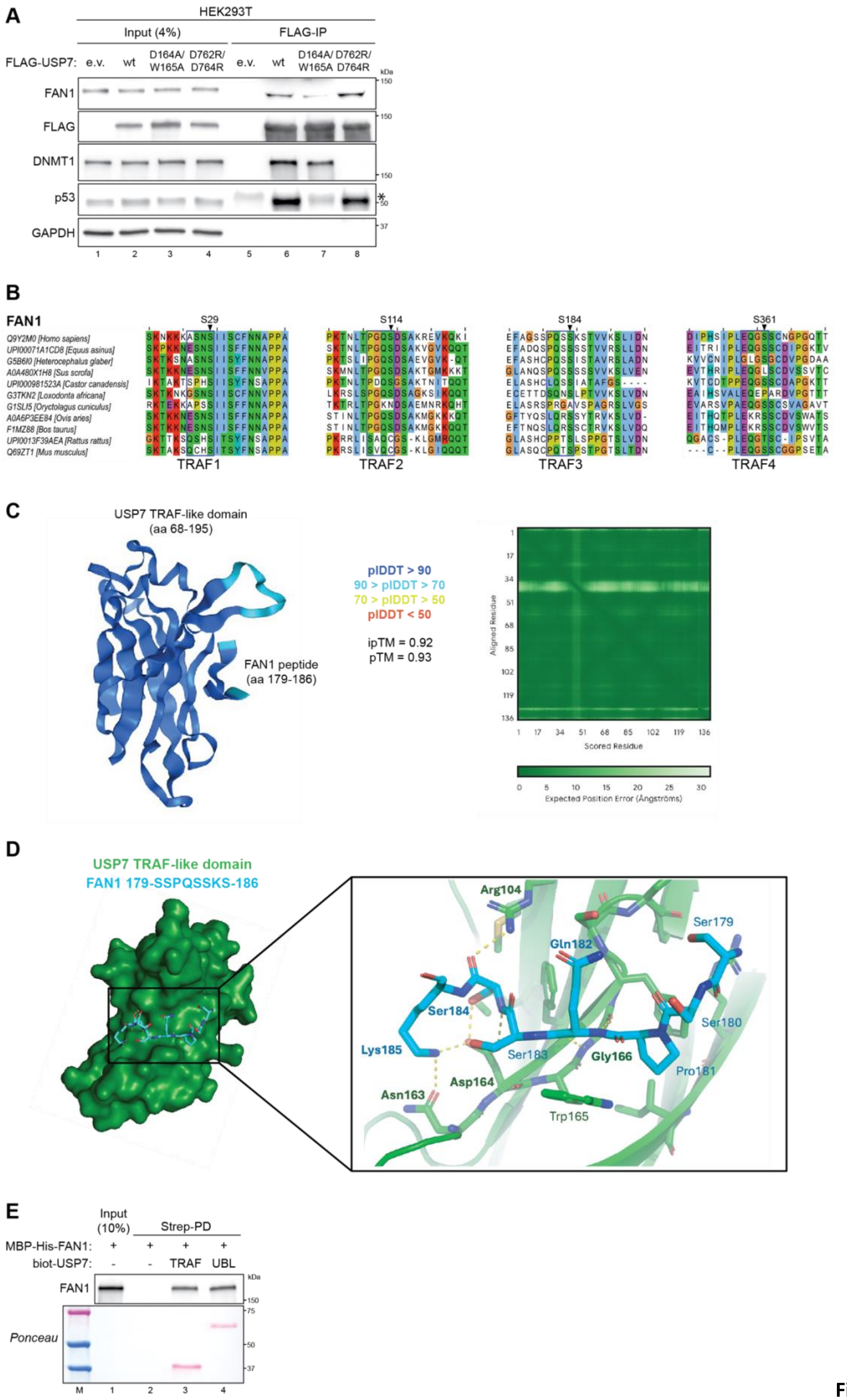
A
HEK293T
Input (4%)
FLAG-IP
FLAG-USP7: e.v. wt D164A/ W165A D762R/ D764R e.v. wt D164A/ W165A D762R/ D764R
FAN1
FLAG
DNMT1
p53
GAPDH
kDa
150
150
150
*
50
37
1 2 3 4 5 6 7 8
B
FAN1
S29
S114
S184
S361
Q9Y2M0 [Homo sapiens]
UPI00071A1CD8 [Equus asinus]
G5B6I0 [Heterocephalus glaber]
A0A480X1H8 [Sus scrofa]
UPI000981523A [Castor canadensis]
G3TKN2 [Loxodonta africana]
G1SLI5 [Oryctolagus cuniculus]
A0A6P3EE84 [Ovis aries]
F1MZ88 [Bos taurus]
UPI0013F39AEA [Rattus rattus]
Q69ZT1 [Mus musculus]
TRAF1
TRAF2
TRAF3
TRAF4
C
USP7 TRAF-like domain
(aa 68-195)
FAN1 peptide
(aa 179-186)
pIDDT > 90
90 > pIDDT > 70
70 > pIDDT > 50
pIDDT < 50
ipTM = 0.92
pTM = 0.93
Aligned Residue
Scored Residue
Expected Position Error (Ångströms)
D
USP7 TRAF-like domain
FAN1 179-SSPQSSKS-186
Arg104
Ser179
Gln182
Ser184
Ser180
Lys185
Ser183
Gly166
Pro181
Asp164
Asn163
Trp165
E
Input (10%)
Strep-PD
MBP-His-FAN1: + + + +
biot-USP7: - - TRAF UBL
FAN1
kDa
150
75
50
37
Ponceau
M 1 2 3 4

Figure

**S2 (related to Figure 2). USP7 contacts FAN1 via both its TRAF-like and UBL domains.**

(**A**) HEK293T cells were transfected with either e.v. or the indicated FLAG-USP7 expression constructs. 48 h later, cells were lysed and whole-cell extracts were subjected to IP using anti-FLAG M2 affinity resin. Inputs and recovered protein complexes were analysed by immunoblotting. Asterisk indicates heavy chain IgGs from anti-FLAG M2 affinity resin. (**B**) Selected fragments from a multiple sequence alignment of human FAN1, focused on four serine residues embedded within the P/A/ExxS motif preferentially recognized by USP7 on various partners, are shown using Jalview[79]. Representative mammalian sequences are included, with the blue box highlighting the positions of the conserved motif surrounding serines S29, S114, S184, and S361. (**C**) Schematic representation of the 3D structure of the interaction complex between USP7's TRAF domain (aa 68-195) and FAN1 peptide (aa 179-186) by AlphaFold 3[41], colored according to the predicted local Distance Difference Test (plDDT). Dark blue: regions with pIDDT greater than 90 (Very high); light blue: regions with pIDDT between 70 and 90 (Confident); yellow regions with pIDDT between 50 and 70 (Low); orange: regions with pIDDT lower than 50 (Very low). (**D**) Analysis of the interactions mediated in the model predicted in **C** indicate S184, as well as other residues in the FAN1 peptide (cyan), may mediate hydrogen bonds with the TRAF domain of USP7 (green). Residues involved in hydrogen bonds are shown in bold. (**E**) Purified recombinant biotin-tagged USP7 fragments (6 µg) were immobilized on Strep-Tactin®XT-4Flow® resin and incubated with purified recombinant MBP-His-FAN1 (1 µg). Ponceau staining was performed to visualize the USP7 fragments. Inputs and recovered MBP-His-FAN1 were analysed by immunoblotting.

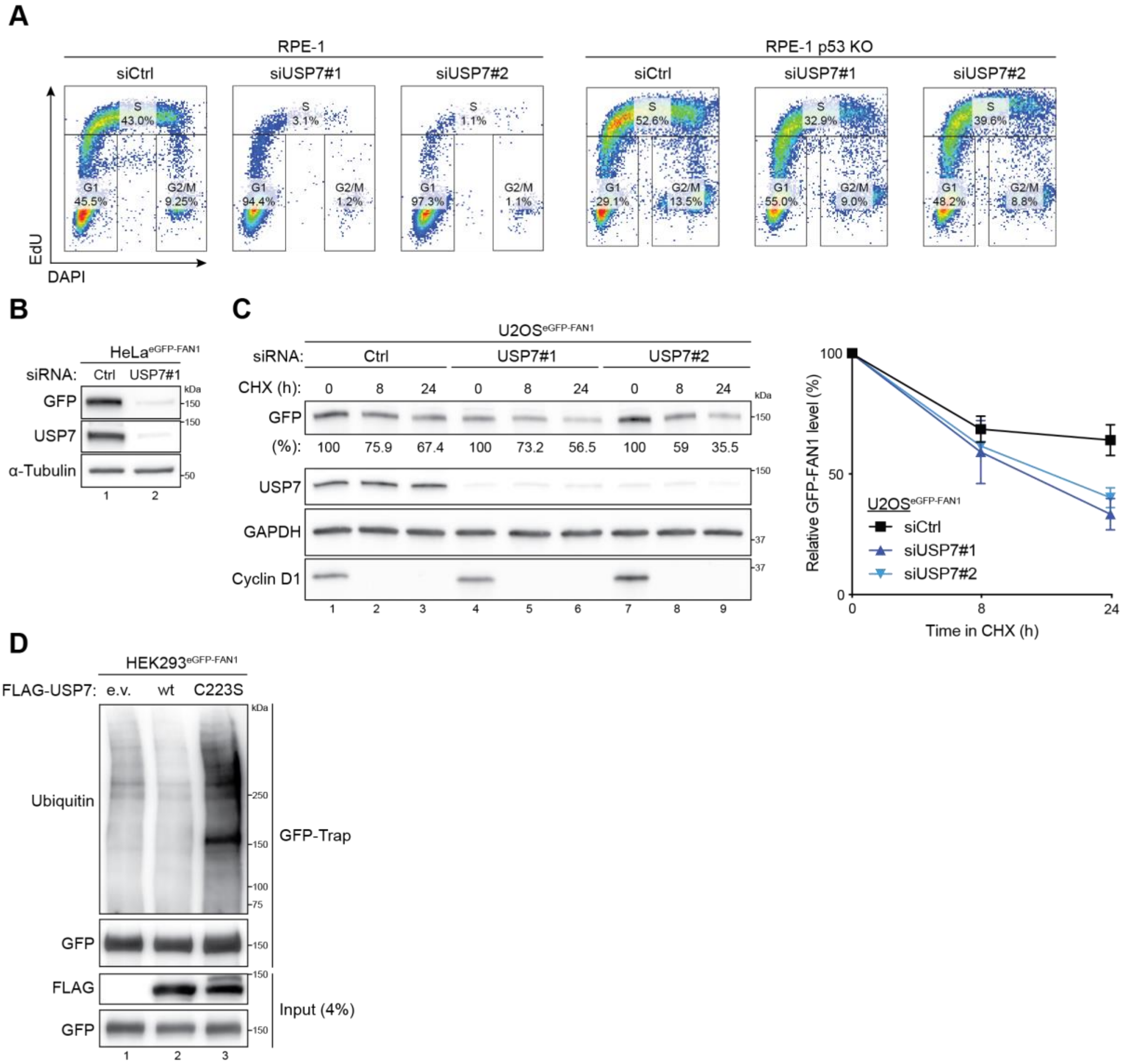


**Figure S3 (Related to Figure 3). USP7 controls FAN1 stability via its deubiquitinase activity.**

(**A**) RPE-1 parental and p53 KO cells were transfected with either non-targeting (Ctrl) or USP7 siRNA oligos. 48 h later EdU was incorporated for 30 min. Cells were harvested, fixed and immunostained with Click-IT reaction and DAPI. Fluorescent intensities were measured by flow cytometry. Dot plots illustrate the EdU signal intensity (y-axis) versus the DAPI content (x-axis). Gates depict the percentage of cells in G1, S and G2/M. (**B**) Doxycycline-inducible HeLa$^{eGFP\text{-}FAN1}$ cells were transfected with either non-targeting (Ctrl) or USP7 siRNA oligos. 24 h later, GFP-FAN1 expression was induced with Dox (100 ng/ml) for 24 h and whole-cell lysates were analysed by immunoblotting. (**C**) Doxycycline-inducible U2OS$^{eGFP\text{-}FAN1}$ cells were transfected with either non-targeting (Ctrl) or USP7 siRNA oligos. 24 h later, GFP-FAN1 expression was induced with Dox (100 ng/ml). 24 h post induction, cells were either mock-treated or treated with CHX (100 µg/ml) for the indicated time points and whole-cell lysates were analysed by immunoblotting (left panel). Relative FAN1 protein levels were determined by quantification of FAN1 band intensity (normalized to GAPDH) with the ImageJ software (right panel). Data are represented as mean values of densitometric quantification ± SD (*n*=3). (**D**) Doxycycline-inducible HEK293$^{eGFP\text{-}FAN1}$ cells were transfected with either e.v. or indicated FLAG-USP7 expression constructs. 24 h later, GFP-FAN1 expression was induced with Dox (100ng/ml). 24 h post induction, cells were lysed and whole-cell extracts were subjected to GFP-Trap resin. Inputs and recovered protein complexes were analysed by immunoblotting.

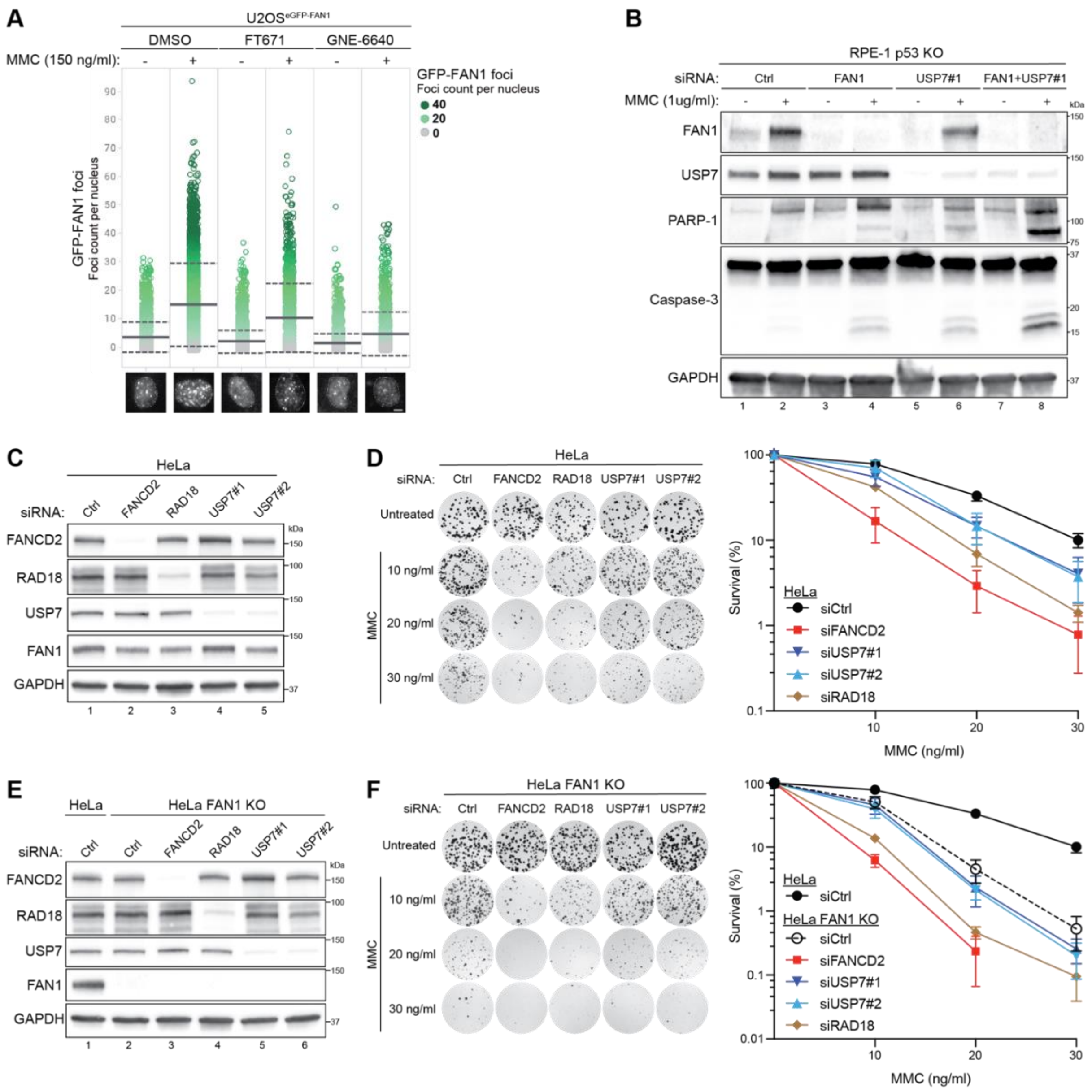

**Figure S4 (Related to Figure 4). USP7 facilitates FAN1 localization and function during ICL repair.**

(**A**) QIBC analysis of chromatin-bound GFP-FAN1 foci in U2OS$^{GFP\text{-}FAN1}$ cells were concurrently mock-treated (-) or treated (+) with MMC (150 ng/ml) and either FT671 (10 µM) or GNE-6640 (10 µM) for 24 h. Color-coded scatterplots indicate the number of GFP-FAN1 foci per nucleus. Mean (solid line) and standard deviation (SD) from the mean (dashed lines) are indicated. Representative images are shown below. Scale bar, 10 µm. (**B**) RPE-1 p53 KO cells were transfected either non-targeting (Ctrl) or USP7 siRNA oligos. 24 h later, cells were either mock-treated (-) or treated (+) with MMC (1 µg/ml) for 48 h and whole-cell lysates were analysed by immunoblotting. (**C**) Western blot analysis of HeLa cells depleted of the indicated factors. (**D**) Clonogenic survival assay of the HeLa cells depleted of the indicated factors and exposed to increasing doses of MMC for 24 h. Viability of untreated cells was defined as 100%. Data are presented as the means ± SD. Representative images are shown. (**E**) Western blot analysis of HeLa and HeLa FAN1 KO cells depleted of the indicated factors. (**F**) Clonogenic survival assay of HeLa FAN1 KO cells depleted of the indicated factors and exposed to increasing doses of MMC for 24 h. Viability of untreated cells was defined as 100%. Data are presented as the means ± SD. Representative images are shown.

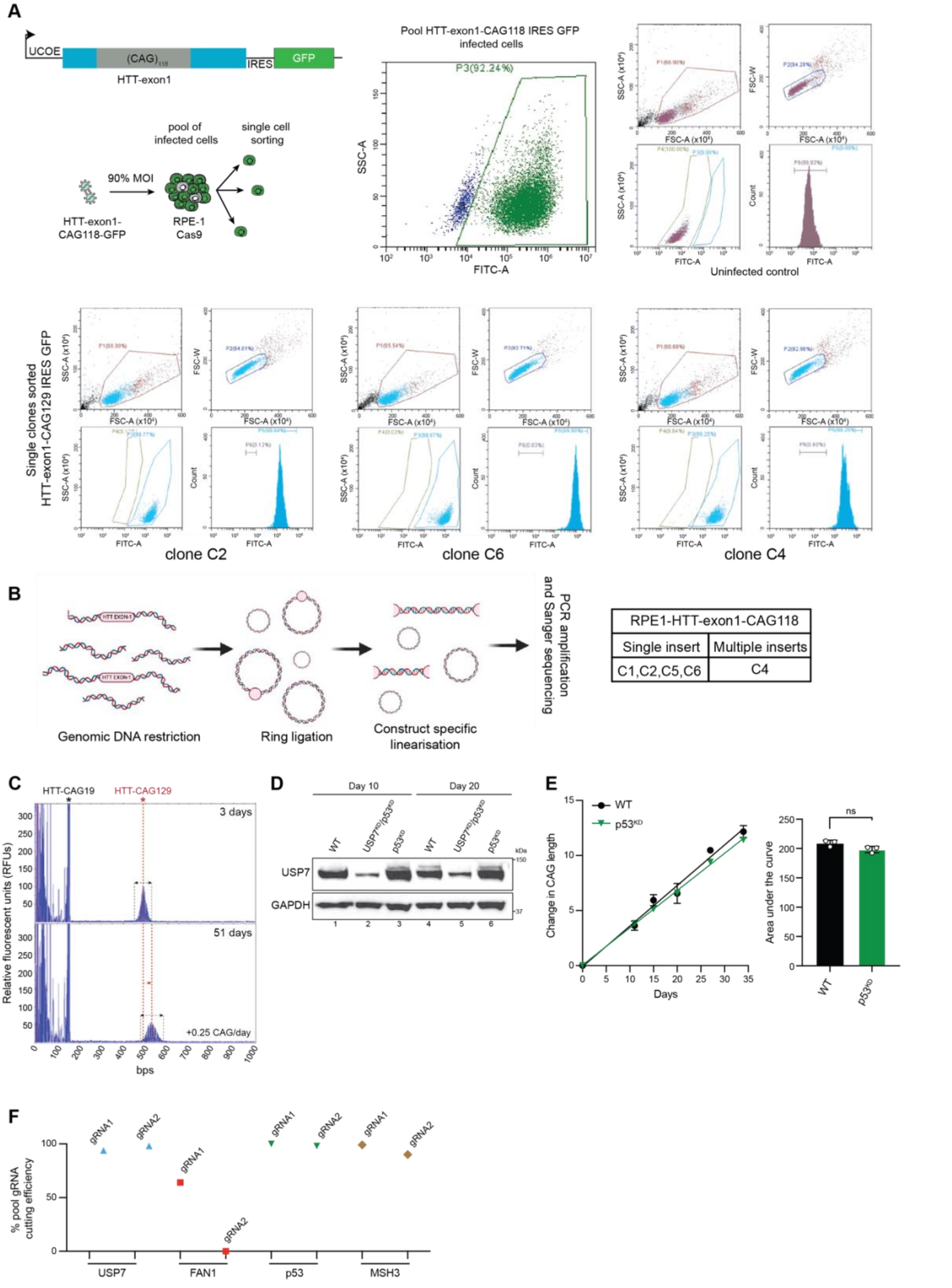


**Figure S5 (Related to Figure 5). The RPE-1 HTT-exon1-CAG129 cell line is a rapid-expansion human model of CAG repeat instability.**

**(A)** Workflow for generating the RPE-1$^{CAG129}$ cell line. *Top:* Schematic of the HTT-exon1-CAG118-IRES-GFP construct containing 118 CAG repeats. RPE-1 Cas9 cells were infected at 90% MOI and single-cell sorted based on GFP fluorescence. FACS plots show GFP intensity in pooled infected cells versus uninfected controls, and in representative clones (C2, C6, C4). By the time of clone selection, sizing and banking, the CAG expanded to 129 repeats. **(B)** Ring ligation strategy to determine genomic integration. Genomic DNA was digested with restriction enzymes, circularized, re-linearized at construct-specific sites, and analysed by PCR and Sanger sequencing. Clones C1, C2, C5, and C6 contained single integrations; clone C4 carried multiple insertions. **(C)** Representative fragment analysis traces showing CAG repeat expansion over time in RPE-1$^{CAG129}$ clone C2. Traces from day 3 and day 51 demonstrate a shift in the HTT-CAG129 peak position, indicating progressive repeat expansion. Peaks correspond to the endogenous HTT-CAG19 allele (black asterisk) and the integrated HTT-CAG129 construct (red asterisk). The expansion rate was estimated at +0.25 CAG repeats per day. Red arrow indicates the broadening of the inserted allele peak over time, reflecting heterogeneity in repeat length, a hallmark of ongoing somatic expansion. (**D**) Western blot validation of USP7 knock-down in RPE-1$^{CAG129}$ cells. Whole cell lysates from wt, USP7 KD, p53 KD, and USP7 KD/p53 KD cells were collected at days 10 and 20 post-infection and analysed by immunoblotting. (**E**) Quantification of sgRNA cutting efficiency in pooled RPE-1$^{CAG129}$ cells following lentiviral-mediated knock-down of USP7, FAN1, p53 and MSH3, as used in experiments shown in Figure 5. **(F)** CAG repeat expansion over time in RPE-1$^{CAG129}$ cells following *p53* knock-down. Left: Time-course analysis of CAG repeat length in *p53*$^{KD}$ (green) and wt (black) cells. Right: AUC quantification shows no significant difference (ns); bars represent mean ± SD (n=3). This experiment was repeated n=3 independent times.

**Supplementary Table 2. List of gRNAs used in this study**

| Gene | gRNA name | Source | 5'-3' sequence | Cell line used |
|---|---|---|---|---|
| *FAN1* | FAN1-466 | This study | CACCACAAGGAAACTCCGA | HeLa |
| *FAN1* | FAN1-470 | This study | AGAAGTGGTTGAAAAACGTG | HeLa |
| *FAN1* | FAN1 gRNA1 | This study | ATTGTCCAGAAAATACGTAA | RPE-1-Cas9-CAG118 |
| *FAN1* | FAN1 gRNA2 | This study | AATGCAGGCTTTCTACAGAC | RPE-1-Cas9-CAG118 |
| *MSH3* | MSH3 gRNA1 | This study | AAATCCACCTCCTCCTCCACAGG | RPE-1-Cas9-CAG118 |
| *MSH3* | MSH3 gRNA2 | This study | CAGAAGAAAGAAGAGACCATTGG | RPE-1-Cas9-CAG118 |
| *TP53* | TP53 gRNA1 | This study | TCTCGAAGCGCTCACGCCCACGG | RPE-1-Cas9-CAG118 |
| *TP53* | TP53 gRNA2 | This study | CCTTGAGTTCCAAGGCCTCATTC | RPE-1-Cas9-CAG118 |
| *USP7* | USP7 gRNA1 | This study | CCACATAATCCACAAATGATTCA | RPE-1-Cas9-CAG118 |
| *USP7* | USP7 gRNA2 | This study | CCGACGTAGCCTGTGTGCTTCTT | RPE-1-Cas9-CAG118 |

**Supplementary Table 3. List of DNA primers used in this study**

| Primer/oligonucleotide | Source | 5'-3' sequence |
|---|---|---|
| FAN1_S184A_F | This study | ACAGAGTGCCAAATCCACAGTTGTTAAGAGCC |
| FAN1_S184A_R | This study | GATTTGGCACTCTGTGGACTAGAACCGG |
| FAN1_1-372_F | This study | CAACCGGTTAACTCGAGTCTAGAGGGCCC |
| FAN1_1-372_R | This study | CGAGTTAACCGGTTGTTTGACCAGG |
| FAN1_del1-372_F | This study | GCCGCATGCCTTACTACCTTCGGAGTTTCCTTG |
| FAN1_del1-372_R | This study | AGTAAGGCATGCGGCCGCTGTACTTG |
| FAN1_del1-40_F | This study | GCCGCATGAAACTTGCCTGCCCCGTTTG |
| FAN1_del1-40_R | This study | CAAGTTTCATGCGGCCGCTGTACTTG |
| FAN1_del157-207_F | This study | GCCTAGCAAACAGTTCTCAAAAAGAAAACGTG |
| FAN1_del157-207_R | This study | AACTGTTTGCTAGGCTTCCCAAACAAATG |
| USP7_TRAF_F | This study | TTGCGTGGTGAGGATCCGGAATTCCGCC |
| USP7_TRAF_R | This study | ATCCTCACCACGCAACTCCATGGGG |
| USP7_delUBL_F | This study | GGCAGGAATGAGGATCCGGAATTCCGCC |
| USP7_delUBL_R | This study | ATCCTCATTCCTGCCGCTCCTTCCG |
| USP7_UBL_F | This study | CCGGTATGGATATTCCTCAGCAGTTGGTGGAG |
| USP7_UBL_R | This study | GAATATCCATACCGGTCTTGTCATCGTC |
| USP7_D164A_W165A_F | This study | AAAATGCGGCGGGATTTTCCAATTTTATGGCCTGG |
| USP7_D164A_W165A_R | This study | ATCCCGCCGCATTTTCTTTATGGAAGAACAAATGAC |
| USP7_C223S_F | This study | AGCGACTAGCTACATGAACAGCCTGCTACAGACG |
| USP7_C223S_R | This study | ATGTAGCTAGTCGCTCCCTGATTCTTTAAGCC |

| | | |
|---|---|---|
| HD3 FAM | Teo *et al.*, 2008 | CCTTCGAGTCCCTCAAGTCCTT |
| HD5 | Mangiarini *et al.*, 1996 | CGGCTGAGGCAGCAGCGGCTGT |

**Supplementary Table 4. List of siRNA oligonucleotides used in this study**

| **Oligonucleotide** | **Sequence** | **Source** | **Concentration used** |
|---|---|---|---|
| siCrtl | CGUACGCGGAAUACUUCGA | Microsynth | 40 nM |
| siFAN1 | GUAAGGCUCUUUCAACGUA | Microsynth<br>MacKay *et al.,* 2010 | 40 nM |
| siFANCD2 | CAGAGUUUGCUUCACUCUCUA | Microsynth<br>Enoiu *et al.,* 2012 | 40 nM |
| siRad18 | AUGGUUGUUGCCCGAGGUU | Microsynth<br>Porro *et al.,* 2017 | 40 nM |
| siUSP7#1 | GUGUAAAGAAGUAGACUAU | Microsynth<br>Jiang *et al.,* 2017 | 40 nM |
| siUSP7#2 | CCCAAAUUAUUCCGCGGCAAA | Microsynth<br>Zlatanou *et al.,* 2016 | 40 nM |
| siUSP9X | CUGUGAUUCAGCAACUCUA | Microsynth<br>Shang *et al.,* 2019 | 40 nM |
| siUSP11 | AAGCGUUACUAUGACGAGGUAUU | Microsynth<br>Wiltshire *et al.,* 2010 | 40 nM |
| siUSP48 | GCGUAAGCAAAGUGUGGAUA | Microsynth<br>Uckelmann *et al.,* 2018 | 40 nM |

**Supplementary Table 5. List of antibodies used in this study**

| **Antibody** | **Species** | **Source** | **Application** | **Dilution** |
|---|---|---|---|---|
| Caspase-3 | Rabbit | Cell Signaling (9662) | IB | 1:1000 |
| Cyclin D1 | Rabbit | Cell Signaling (2922) | IB | 1:1000 |
| DNMT1 | Rabbit | Cell Signaling (5032) | IB | 1:1000 |
| FAN1 | Rabbit | ProteinTech (17600-1-AP) | IB | 1:1000 |
| FAN1 | Mouse | Genscript (1A11-2-A) | IB and *PLA* | 1:500<br>*1:100* |
| FAN1 | Rabbit | Novus (NBP1-42677) | IP | 1:1000 |
| FAN1 | Goat | MacKay et al. (2010) | IF | 1:200 |
| FANCD2 | Rabbit | Novus (NB-100-182) | IB and *IF* | 1:5000<br>*1:250* |
| FLAG M2 | Mouse | Sigma-Aldrich (#F1804) | IB | 1:1000 |
| GAPDH | Mouse | Millipore (MAB374) | IB | 1:40000 |
| GFP | Rabbit | Abcam (ab290) | IB | 1:1000 |
| GFP | Mouse | Roche (11814460001) | IF | 1:200 |
| GST | Rabbit | Abcam (ab9085) | IB | 1:500 |

| | | | | |
|---|---|---|---|---|
| HA | Mouse | Santa Cruz (sc-7392) | IB | 1:1000 |
| Lamin B1 | Rabbit | Abcam (#ab16048) | IB | 1:1000 |
| MLH1 | Rabbit | Abcam (ab92312) | IB | 1:5000 |
| p53 | Mouse | Santa Cruz (sc-126) | IB | 1:4000 |
| PARP-1 | Rabbit | Abcam (ab227244) | IB | 1:1000 |
| RAD18 | Rabbit | Cell Signaling (9040) | IB | 1:1000 |
| RNF169 | Rabbit | Aviva Systems Biology Corp (ARP43508_P050) | IB | 1:1000 |
| Ubiquitin | Mouse | Santa Cruz (sc-8017) | IB | 1:1000 |
| USP7 | Rabbit | Bethyl (A300-033A) | IB | 1:10000 |
| USP9X | Rabbit | Bethyl (A301-351A-T) | IB | 1:2000 |
| USP11 | Rabbit | Bethyl (A301-613A) | IB | 1:2000 |
| USP48 | Rabbit | Abcam (ab72226) | IB | 1:2000 |
| α-Tubulin | Mouse | Sigma-Aldrich (#T9026) | IB | 1:20000 |

**Supplementary Table 6. List of peptides used in this study**

| **Peptide** | **Sequence** | **Source** |
|---|---|---|
| TRAF1 | biotin-NKKKASNSIISV-NH2 | This study (SynPeptide) |
| TRAF2 | biotin-TNLTPGQSDSAK-NH2 | This study (SynPeptide) |
| TRAF3 | biotin-AGSSPQSSKSTV-NH2 | This study (SynPeptide) |
| TRAF3 mut | biotin-AGSSPQSAKSTV-NH2 | This study (SynPeptide) |
| TRAF4 | biotin-PLEQGSSCNGPG-NH2 | This study (SynPeptide) |
| UBL1 | biotin-SDSAKREVKQKIS-NH2 | This study (SynPeptide) |
| UBL2 | biotin-SLASKLSRKYVKA-NH2 | This study (SynPeptide) |
| UBL3 | biotin-VKAKKSIDKDEEF-NH2 | This study (SynPeptide) |
| FAN1 60mer | Human FAN1 (118-177) | Porro *et al.*, 2021 (GenScript) |
